# Primitives for motion segmentation in the retina

**DOI:** 10.64898/2025.12.31.697115

**Authors:** Samuele Virgili, Thomas Buffet, Matthew Chalk, Olivier Marre

## Abstract

The center surround structure of ganglion cells receptive fields is suited for edge detection, a first step towards image segmentation. However, in a dynamical visual scene with moving, textured objects, it is less clear how the retina represents the local boundaries of these objects. Here, we show that the spatial selectivity of ganglion cells changes during their responses to a moving object. Specific cell types only respond to contrast changes near the edges of the moving object, while being insensitive to changes in other parts of their receptive field. Using a non-linear model to reproduce this result, we could isolate the mechanism responsible for this selective representation. These types of ganglion cells represent selectively textures only when they are near moving edges, and may thus provide useful primitives for motion segmentation.

## Introduction

The visual system converts patterns of light into structured representations of objects and surfaces. A central step in this transformation is image segmentation ^1^, the process by which a static image is parsed into meaningful components that form the basis of our perceptual interpretation of the scene. A prerequisite for segmenting the visual world is the extraction of low-level features that can be grouped into distinct objects. The retina contributes critically to this early processing stage, most notably through mechanisms that enhance edges and suppress uniform regions. The receptive field of most retinal ganglion cells (RGCs) ^2,3^ has a center–surround structure that amplifies responses to edges while attenuating responses to homogenous areas. While image segmentation is not performed in the retina, this suggests that the retina provides primitives for this task, i.e. computations whose output can be useful for this. For static images, the retina thus provides a primitive for image segmentation by enhancing edges.^4,5^

Natural scenes, however, rarely remain static. To parse a dynamic visual scene into independently moving objects, the brain must detect the moving boundaries of these objects. How retinal computations may contribute to this process, known as motion segmentation, remains much less clear. It is unclear whether the retina can provide useful low-level features for motion segmentation. In particular, can the retina selectively emphasize the local texture boundaries in moving objects?

Although retinal responses to moving stimuli have been studied extensively, previous works have largely focused on simple objects and on mechanisms for encoding motion direction ^6,7^, tracking object position ^8–10^, or detecting motion ^11,12^. Less is known about how the retina encodes moving objects with complex spatial content, such as textured surfaces. Tackling this question is necessary to understand the contribution of the retina to motion segmentation.

Here, we investigate the spatial selectivity of RGCs during stimulation with moving textured objects. To this end, we developed a perturbative stimulation approach that allows us to probe RGC selectivity across space and time when RGCs respond to a moving object. We find that, for specific RGC types, spatial selectivity shifts over time as an object moves: they only detect contrast changes near the object’s boundary. Distinct RGC types preferentially encode different portions of a moving object, with some tracking the texture content near the leading edge and others the one of the trailing edge. Spatial selectivity is therefore not “static”, but reshaped by the presence of moving objects. We show that these behaviors can be reproduced by a nonlinear model, and that the subunit rectification of this model is essential for a robust “edge tracking” selectivity. These results show that different RGC types provide complementary representation of the texture of moving objects, with specific types selectively representing the texture near the edge of moving objects. They may supply key primitives for motion segmentation at the earliest stages of visual processing.

## Results

Objects in motion are not defined solely by their trajectory or speed, they also have a texture and shape, features that are essential for object identification. RGCs need to convey information about these features to downstream areas of the brain, but the way they do so is still unclear. Here we asked if ganglion cells are sensitive to the texture of an object when this object is moving. To address this, we measured their response to different texture manipulations superposed on top of a moving object.

### Controlled contrast perturbations reveal that some ganglion cells are selectively tuned to the leading edge of moving textures

We first presented mouse retinal explants with a uniform bright bar moving at constant speed (Figure 1A) and recorded RGC responses using a multi-electrode array (see Methods). To test whether the ganglion cells are selective to changes in the texture, we selectively increased the contrast of either the leading edge (Figure 1B, orange) or the trailing edge (Figure 1B, green) of the same bar. Different RGC types responded qualitatively differently to the three stimuli. Interestingly, some cells changed their response compared to the uniform case only when the leading edge of the bar was enhanced, remaining nearly unaffected by the trailing edge contrast increase (Figure 1 C,D, right). This was the case of the ‘ON high frequency’ and ‘ON transient DS’ RGC types, as defined in ^13^. Other cells changed their response for both contrast modifications, showing increased firing early when the leading edge was enhanced and later when the trailing edge was (Figure 1 C,D, left). These differences suggest that some RGC types are only sensitive to contrast variations near the leading edge of a moving object and are invariant to contrast variations close to the trailing edge, while others have a behavior more consistent with spatial encoding and are sensitive to contrast variations throughout the object.

**Figure 1.**
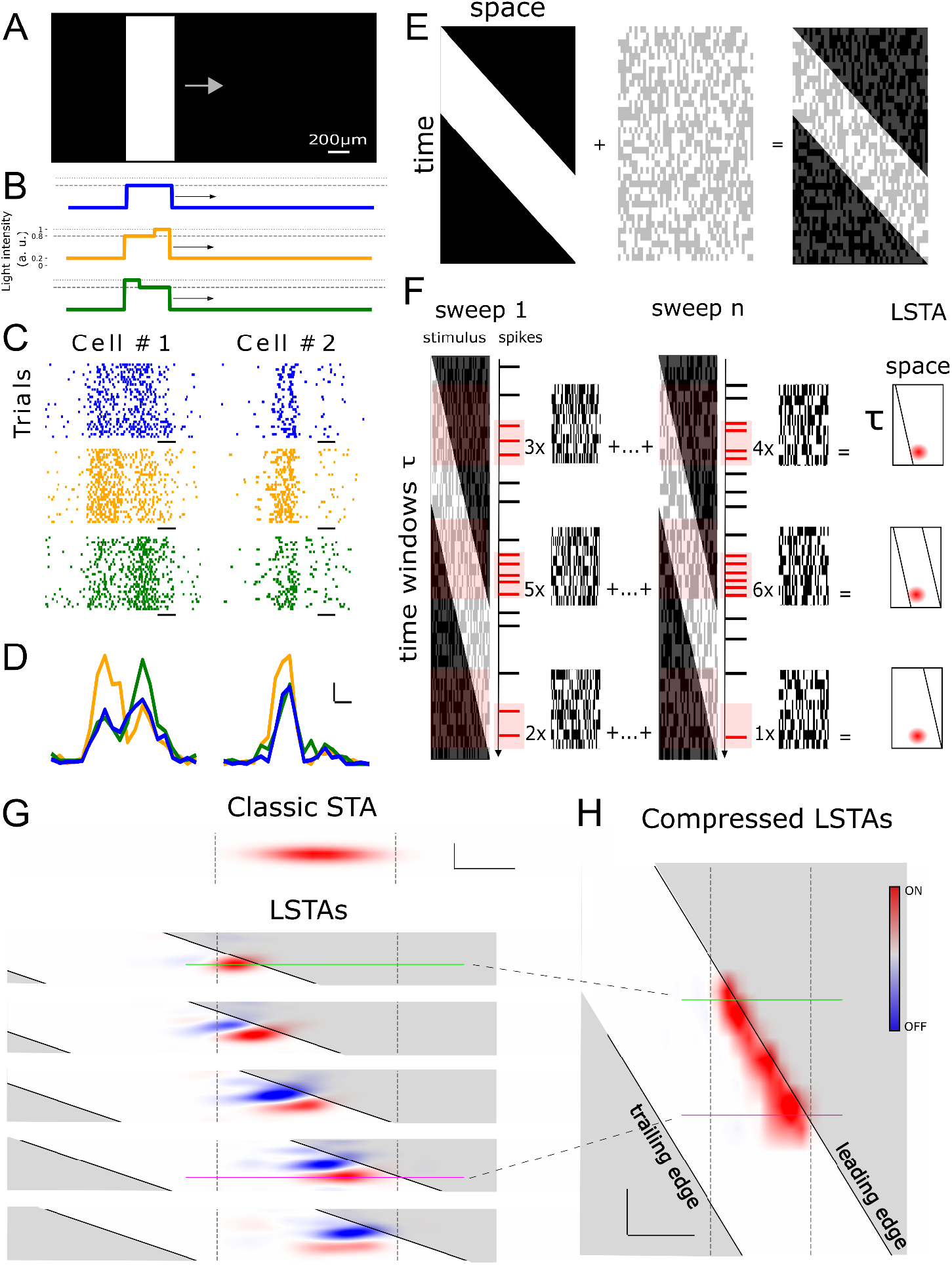
Controlled contrast perturbations reveal that some ganglion cells are selective only to the leading edge of moving textures. **A.** Schematic of the visual stimulus: a uniform bright bar moves at constant speed over a dark background (bar with 0.6 contrast on top of a 0.2 background). Arrow indicates direction of motion. **B**. Luminance profiles of three different textured bars. The uniform bar (blue) is modified through an increase of contrast of either the leading (orange) or the trailing (green) edge. **C**. Raster plots showing the response of two example cells (left and right columns) to the three stimuli shown in B. Each row is a repetition of the same stimulus. Cell #1 responds to both contrast manipulations. Cell #2 responds selectively to leading edge enhancement. Black scale bars are 200 ms. **D**. Peri-Stimulus Time Histograms (PSTHs) of the same two example cells shown in C. Responses are averaged over 20 repetitions and superimposed for comparison. Scale bars are 200 ms along x and 20 Hz along y. **E**. Perturbative stimulation paradigm: The same bar in A (represented in x-t coordinates) is perturbed with random patterns of dim 1D white noise. See Supplementary video 1 for an example perturbed bar sweep. **F**. Schematic of local spike-triggered averages computation. LSTAs were computed in short time windows of size τ (pink boxes) throughout the bar sweep. For each window and each cell, perturbation patterns were averaged across trials, weighted by the number of spikes they evoked. **G**. Classical STA vs LSTAs. Top: The classical STA for an example ON cell obtained by reverse correlating full contrast 1D white noise. The vertical dashed lines highlight the extension in space of the classical RF. Bottom: LSTAs for the same cell obtained for successive positions of the bar. In each filter, the x-axis is space and the y-axis is time lag from spike. The two solid black lines trace the position of the bar edges in space-time. Scale bars are 100 µm along x and 150 ms along y. **H**. Compressed LSTA representation: 1D slices of each LSTA (taken at fixed time lag, magenta and green lines) are stacked over time to highlight the evolution of the cell’s spatial selectivity. This RGC responds selectively to ON contrast at the leading edge of the bar and shows little to no modulation from texture changes elsewhere. The x-axis is space, y-axis is time like in E. Scale bars are 200 µm along x and 100 ms along y.

To understand the origin of these different sensitivities to texture changes, we superimposed low-contrast, one-dimensional white noise patterns on top of a uniform moving bar ^14^(Figure 1E, Supplementary video 1). The contrast of the random patterns was chosen to evoke a small but visible change in the average ganglion cell response compared to the response to the uniform bar (see Methods and Supplementary Fig. 1). We repeated the same bar sweep 2050 times, superimposing the bar each time with a different random pattern (Figure 1F). We then chose several time windows across the bar sweep (pink shaded areas in Figure 1F) corresponding to different bar positions, and for each of them we estimated a Local Spike-Triggered Average (LSTA), as the average of the perturbation patterns presented in that time window across different bar sweeps, weighted by the number of spikes they evoked (Figure 1F).

The LSTAs defined here are similar to a Spike-Triggered Average (STA)^15^ (Figure 1G, top row) in principle, but with an important difference: since the random patterns used to estimate the LSTAs have small contrast, they explore a small, local region of the stimulus space. Unlike classical STAs, which reflect responses to noise alone, our LSTAs capture how cells respond to local contrast variations on top of a strong driving signal, the moving bar. Measuring the LSTAs and their evolution in time, we could thus measure how the spatial selectivity of a cell evolves in time due to the presence of the moving bar and how this spatial selectivity differs from the classically defined receptive field.

We first focused on ganglion cells that belong to the type ‘ON high frequency’ according to the classification proposed in ^13^, which we found above to change their responses only when the leading edge of the bar was changed (Figure 1D). We observed that their LSTAs are smaller than the classical STA, and that their positions change within the receptive field to follow the leading edge of the bar (Figure 1G). To better visualize this change over time, we extracted from each LSTA a 1D slice at fixed time lag (see Methods) and stacked them chronologically (Figure 1H). This representation confirms that the cell spatial selectivity tracked the bar’s leading edge throughout its trajectory. All the cells of this ‘ON high frequency’ type show the same behaviour, with the LSTAs following the leading edge of the bar.

Our LSTA analysis thus shows that for all the duration of the bar sweep across the receptive field, the response of the ‘ON high frequency’ cells was enhanced only by ON changes of contrast on the leading edge of the bar and not across the rest of its surface. This confirmed and expands our finding of a stronger sensitivity for high contrast at the leading edge found in (Figure 1D, right). Thus, when stimulated with a moving texture, this cell type is spatially selective to its leading edge and invariant to the rest. This is at odds with the classical, static, receptive field definition that predicts that a cell should be selective to the same region of space, regardless of the position of the moving object.

### Different retinal ganglion cell types show complementary spatiotemporal selectivity to moving textures

We next asked whether this form of dynamic spatial selectivity extends to other RGC types, and if so, whether some exhibit complementary tuning to other regions of the moving object.

To address this, we recorded the responses of 759 RGCs across 3 mouse retina explants using the perturbed moving bar stimulus (Figure 2A, left; see Figure 1E). From this population, 325 cells (43%) exhibited LSTAs that could be reliably estimated. We functionally classified the recorded cells using a chirp stimulus (Figure 2A, right) and drifting gratings, following the criteria described in ^13^, and focused our analysis on ON-only and OFF-only cell types, excluding ON-OFF types for clarity. We identified 11 cell types with well-defined LSTAs (Figure 2C). We observed several distinct LSTAs modulations across types.

**Figure 2:**
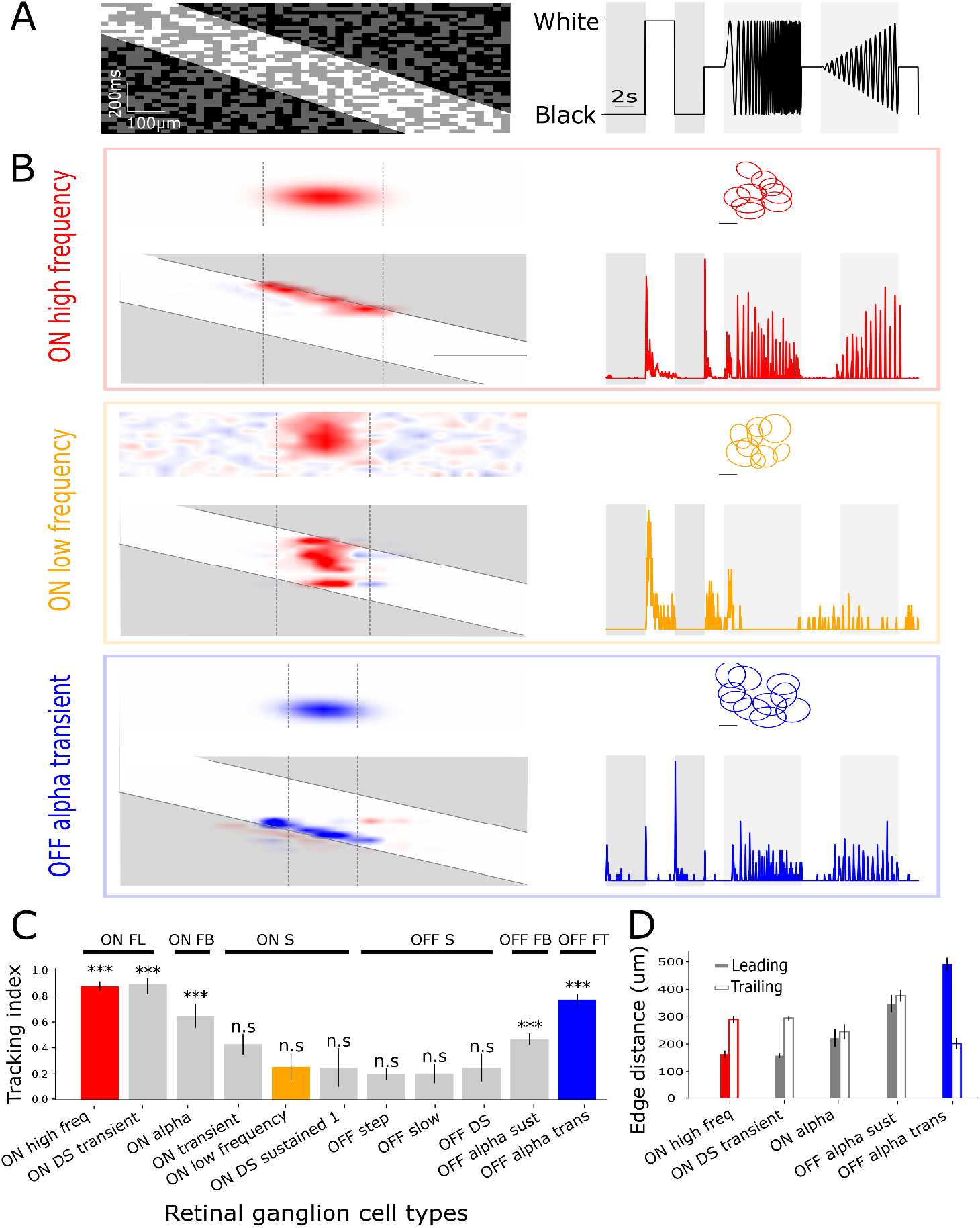
Distinct retinal ganglion cell types show complementary spatiotemporal selectivity to moving textures. **A**. Left: Example of a perturbed moving bar (as in Figure 1E). Right: Temporal profile of the “chirp” stimulus used for functional classification of RGCs^13^. **B**. Representative examples of three identified LSTA behaviors. Top: An ON high frequency cell exhibits LSTAs with ON polarity that track the leading edge of the moving bar (left); the type’s cells mosaic (middle) and the example cell chirp response (right) are shown. Middle: An ON low frequency cell displays static ON LSTAs that remain fixed within the receptive field, consistent with classical luminance encoding. Bottom: An OFF alpha transient cell shows LSTAs with OFF polarity that track the trailing edge of the bar. **C**. Tracking index across all recorded RGC types (n = 11), quantifying how well the spatial position of LSTAs tracks the bar’s leading or trailing edge over time. Tracking was considered significant for ^−4^ Kolmogorov-Smirnoff test, see Methods). Error bars are S.E.M calculated across cells within a type. Based on this index and the distances in D, cell types could be separated in 6 LSTA behaviors: ON following leading edge (ON FL), ON following both edge (ON FB), ON static (ON S), OFF static (OFF S), OFF following both edges (OFF FB) and OFF following trailing edge (OFF FT), see Supplementary Figure 3. **D**. Average distance between each LSTA and the leading or trailing edge of the bar, computed only for types with significant tracking.

As shown in Figure 1, ON high frequency RGCs had ON-polarity LSTAs that tracked the leading edge of the moving bar (Figure 2B, top row). This ‘ON leading edge–tracking’ behavior was also observed in other types, including ON DS transient and ON alpha RGCs. In ON alpha cells, LSTAs tracked both the leading and trailing edges of the bar (Supplementary Figure 2).

To quantify whether a given RGC type’s LSTAs tracked the position of the moving bar, we defined a tracking index (see Methods), measuring how much of the variance in the LSTAs positions over time can be explained by the position of the bar. This index ranges from 0 (static LSTAs) to 1 (perfect tracking of the bar edge). We compared the distribution of tracking indices for each type to a null distribution generated by randomly shuffling LSTA timepoints, and considered a type as ‘tracking’ only if the observed and null distributions were significantly different (*p* < 10^−4^ Kolmogorov-Smirnoff test, Figure 2C). This confirmed that ON high frequency, ON DS transient and ON alpha had a spatial selectivity that tracked the leading edge of the moving bar.

3 other ON types (ON low frequency, ON DS sustained, and ON transient) did not exhibit edge-tracking. Their LSTAs had ON polarity but remained static across time (Figure 2B, middle). These ‘ON static’ cells responded to texture perturbations across the entire classical receptive field, independent of the position of the bar.

Among OFF cell types, two (OFF alpha transient and OFF alpha sustained) showed edge-tracking, with LSTAs that followed the trailing edge of the bar (Figure 2B, bottom). These cells selectively encoded OFF contrast near the rear of the moving object and were insensitive to perturbations on the leading part. As in the ON alpha case, OFF alpha sustained cells also tracked both edges of the bar (Supplementary Figure 3). The remaining OFF types (OFF step, OFF slow, and “OFF DS”) had static LSTAs with OFF polarity that spanned the RF, but were suppressed when the bright bar overlapped the RF, again consistent with classical OFF responses (Supplementary Figure 3).

While the tracking index quantified whether LSTAs followed the bar, it did not specify which edge was being tracked. To address this, for the types identified as tracking, we computed the average distance between the LSTA center and each bar edge (Figure 2D). ON LSTAs were closer to the leading edge, whereas OFF LSTAs were closer to the trailing edge, except for ON and OFF alpha sustained cells, which showed roughly symmetric distances to both edges—consistent with them tracking the two edges.

In summary, we found that cells of the same functional type tend to exhibit consistent LSTA dynamics. We could categorize the LSTA dynamics in 4 categories:

1. ON leading edge–tracking (Figure 2B, top),
2. ON static (Figure 2B, middle),
3. OFF trailing edge–tracking (Figure 2B, bottom),
4. OFF static (Supplementary Figure 3).

Note that several types can share similar LSTA dynamics. These complementary dynamics suggest that RGC types encode different components of a moving object’s texture.

### A subunit model with gain control explains diverse spatial selectivities in ganglion cells during motion

We next investigated which models could reproduce the distinct spatiotemporal selectivity patterns revealed by our LSTA analysis. As argued previously^14^, a LN model cannot account for LSTAs that change position over time. For such models the LSTA would remain unchanged, up to a constant global factor, independent of the background motion stimulus. We thus tested if non-linear models could reproduce our results.

To address this, we modeled ganglion cell processing with a two-stage LNLN (linear–nonlinear–linear–nonlinear, Figure 3A) model ^14,16–21^. The visual stimulus is first convolved with spatiotemporal sub-units that model the photoreceptor to bipolar cell synapses. The output of the convolution is then rectified to account for the bipolar to ganglion cell synaptic transmission. Finally, the output of many subunits is spatiotemporally pooled and again rectified to predict the recorded ganglion cell membrane potential and its spike threshold.

**Figure 3.**
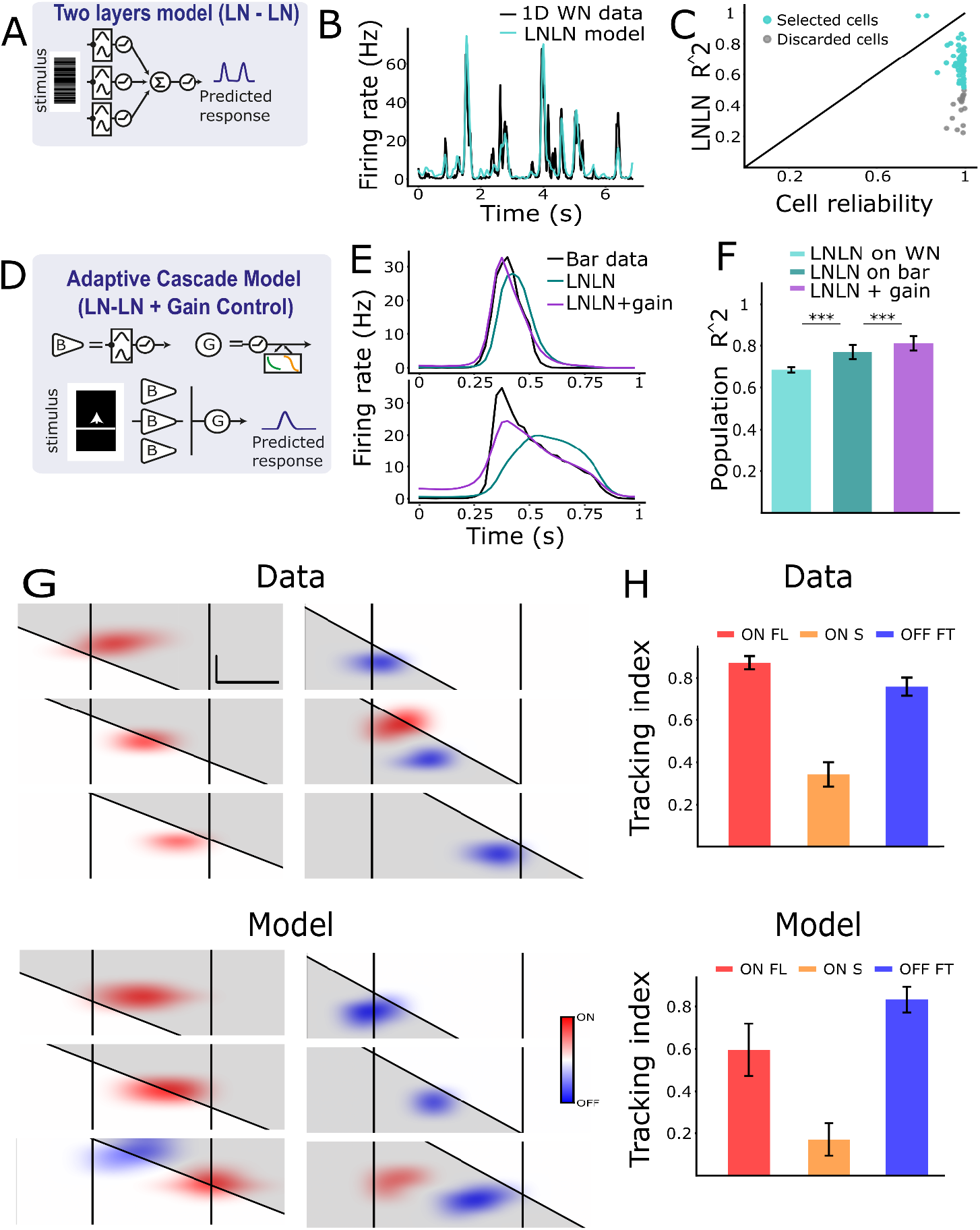
A single subunit model with gain control predicts diverse RGC responses and spatiotemporal selectivity during motion. **A.** Schematic of the two-stage LNLN model architecture. The stimulus is convolved with spatiotemporal subunits, rectified, spatially pooled, and passed through a second rectification to generate the predicted ganglion cell response. **B**. Example of model performance for an example RGC in response to 1D white noise. **C**. Comparison of model performance with response reliability across all modelled cells. Cells with reliable white noise responses (correlation between odd and even trials r > 0.8) and high model fit (R^2^ > 0.5) were selected for downstream analysis (cyan). **D**. Schematic of the extended model including a gain control stage. A divisive normalization term was added after the final nonlinearity to account for adaptation effects. **E**. Firing rate responses to the moving bar for two example cells. The original LNLN model (cyan) misses the onset timing, while the extended model with gain control (purple) corrects the responses onset and the tuning curves shape. **F**. Population performance of the three model variants: trained on white noise, evaluated on the bar without gain control, and with gain control. Gain control significantly improves prediction accuracy (Wilcoxon-signed rank test, *p* < 1*e* ^−8^). **G**. LSTA from perturbation data (top) and model (bottom) for two example cells: ON leading-edge tracking (left) and OFF trailing-edge tracking (right). The model reproduces the spatiotemporal trajectory of LSTAs. Scale bars are 100 µm along x and 150 ms along y. **H**. Data vs model tracking index. Top: Average LSTAs tracking index across recorded cells that showed chirp types linked to either ON following leading edge (ON FL), ON static (ON S) or OFF following trailing edge (OFF FT) behaviors. See Figure 2C for the type’s detail. Bottom: Average index across modelled cells that showed matched chirp types.

The subunit filters were derived from experimentally measured bipolar cell receptive fields^22^. We trained the remaining model parameters (the two rectification thresholds and the pooling weights) on ganglion cell responses to 1D white noise stimuli similar to the ones used in ^22^. We modelled 79 cells across 4 retinae. Cells were selected based on high reliability in their responses to repeated white noise stimuli (correlation across odd and even trials r > 0.8) and clear classification into one of the LSTA behaviors described in Figure 2. For 60 of these cells (76%), the model achieved high performance on held-out white noise data (noise-corrected R^2^ > 0.5; Figure 3B–C) and was used for subsequent analysis.

We then used the model to predict the responses of the selected cells to a uniform moving bar identical to the one used for LSTA estimation, but without added perturbations (Figure 1E, left). For many cells, the model accurately predicted responses to the bar, but for others, it failed to reproduce the correct timing of response onset (Figure 3E, cyan). This is likely due to the fact that since the model was trained on spatiotemporally uncorrelated noise, its parameters failed to account for the gain control mechanisms that have been shown to take place in the retina under stimuli like moving bars ^8,19^.

To address this, we introduced a gain control stage to the model, positioned after the second nonlinearity (Figure 3D). This adaptive mechanism adjusts the model’s output based on recent response history ^8,19^. Adding gain control successfully corrected the timing of the predicted responses (Figure 3E, purple) and led to a significant improvement in prediction accuracy across the population (Figure 3F, Wilcoxon-signed rank test, *p* < 1*e*^−8^) without requiring a full retraining of all the other parameters of the original model (see Discussion).

We next asked if the model could also reproduce the spatiotemporal dynamics of ganglion cell selectivity as measured by the LSTAs. For each cell, we predicted LSTAs from the model using the gradient of the predicted response to the unperturbed bar (see Methods). While the model was not trained on ganglion cell responses to the perturbed movie, it accurately recapitulated the motion-dependent LSTA behavior observed in data, including leading-edge tracking in ON cells and trailing-edge tracking in OFF cells (Figure 3G). To quantify this, we grouped RGCs by their LSTA behaviors (ON following leading edge (ON FL), ON static (ON S), and OFF following trailing edge (OFF FT) as defined by their chirp responses, Figure 2C) and we computed the tracking index introduced in Figure 2 for the model predicted LSTAs. The model’s predictions closely matched experimental values across RGC types (Figure 3H).

These results demonstrate that a single, biologically inspired model architecture comprising rectified spatiotemporal subunits and a gain control mechanism can account for both the firing rate responses and the diversity of RGCs spatial selectivity during motion in a type-specific manner.

This suggests that much of the heterogeneity in motion encoding and LSTA dynamics among RGC types may emerge from a shared computational scaffold shaped by subunit integration and local adaptation dynamics, but where different types have different parameter values. We then asked which parameter values are important to qualitatively change the LSTA dynamics.

### Subunit rectification threshold controls edge-tracking versus static selectivity

Having established that a single model architecture can account for the distinct LSTA behaviors observed across RGC types, we next asked: what are the key internal parameters that determine whether a modeled cell exhibits edge-tracking LSTAs (i.e., sensitivity to contrast modulations near the moving object’s leading or trailing edge) or static LSTAs (i.e., sensitivity across a fixed spatial region of the receptive field)?

We focused on three parameters that could shape the model’s spatiotemporal selectivity: (1) the subunit rectification thresholds, which control the level of input required to activate a subunit; (2) the readout threshold, applied after spatiotemporal pooling across subunits; and (3) the strength of the gain control, which modulates responses based on recent activity history.

To identify which of these parameters most strongly affected LSTA behavior, we considered cells from 3 different LSTA behaviors (ON following leading edge (ON FL), ON static (ON S), and OFF following trailing edge (OFF FT), like in Figure 3H) and compared the mean values of each parameter across modelled cells in these groups (Figures 4B–D). Only the subunit threshold (Figure 4B) showed a consistent relationship with LSTA behaviour: cells with edge-tracking LSTAs (ON FL, OFF FT) had significantly higher thresholds than cells with static LSTAs (ON S). Detailed data in Supplementary Table 1. In contrast, neither the readout threshold (Figure 4C) nor gain control strength (Figure 4D) reliably differentiated between the groups.

**Figure 4.**
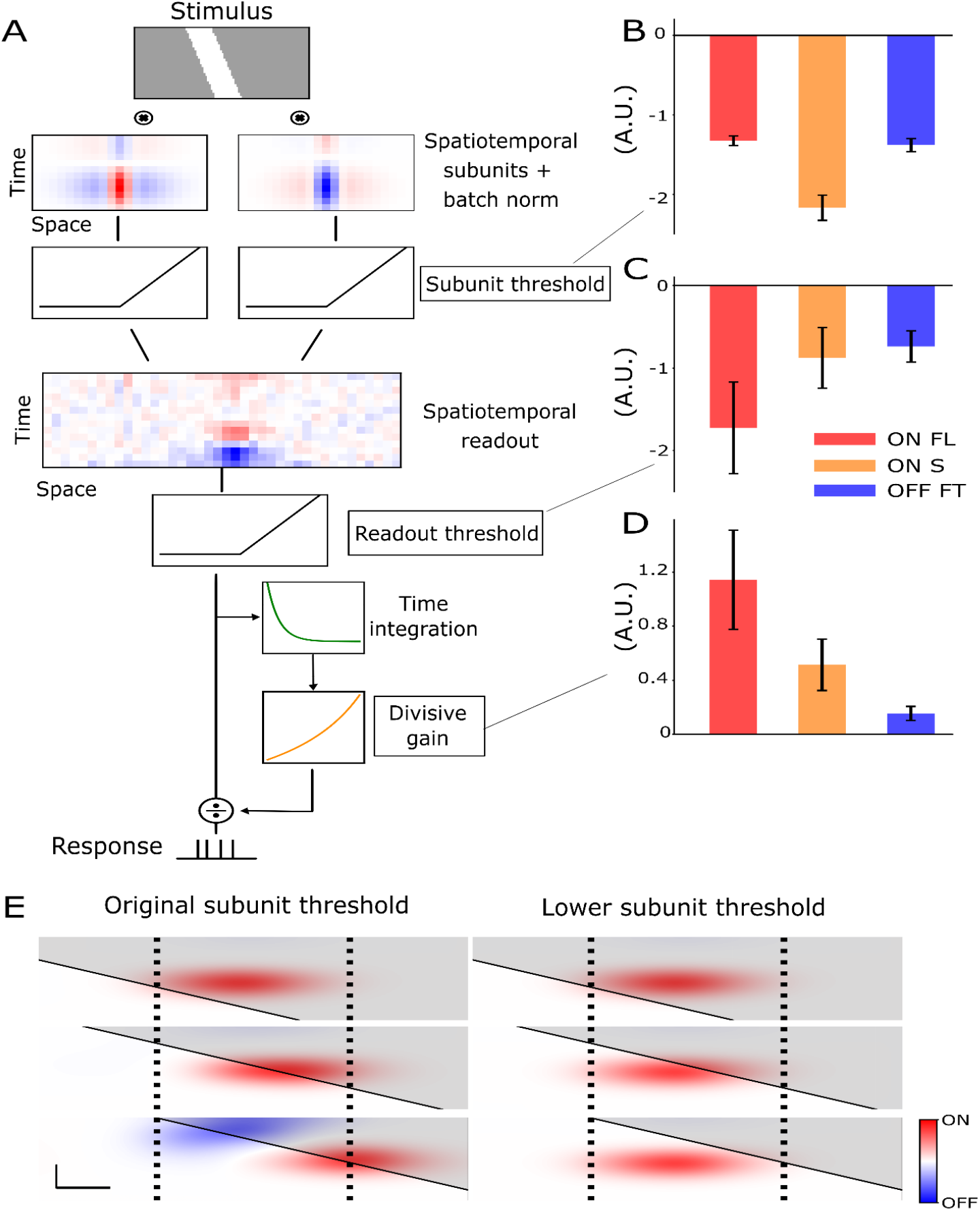
Subunit rectification threshold controls edge-tracking versus static selectivity in ganglion cell models. **A.**Schematic of the full model architecture. A moving bar stimulus is convolved with two shared spatiotemporal subunits (one ON, one OFF), followed by batch normalization and rectification. Subunit outputs are pooled in space and time and passed through a second rectification, then integrated over time and modulated by a divisive gain control mechanism to generate the predicted firing rate. **B–D**. Comparison of fitted model parameters across cells grouped by LSTA behavior: ON following leading edge (ON FL, red), ON static (ON S, orange), and OFF following trailing edge (OFF FT, blue). **B**. parameters comparison for the subunit threshold. Because of the batch-normalization layer, the subunit activations range from -5 to 5. The values reported here are from the OFF model subunit, as almost all modelled cells relied exclusively on this subunit for their predictions. **C**. parameters comparison for the readout threshold. **D**. parameters comparison for the gain control strength. Table of significance testing across conditions in Supplementary Table 1. **E**. Model-predicted LSTAs for one example cell before (left) and after (right) lowering the subunit rectification threshold. Lowering the threshold causes the LSTAs, which originally tracked the bar edge, to become spatially fixed—switching from edge-tracking to static behavior. Dashed lines denote the spatial extent of the cell’s classical RF. Scale bars are 100 µm along x and 150 ms along y.

To directly test the specific effect of subunit threshold on LSTA behavior, we used the model to simulate how the LSTAs of an ON FL cell would change if we lowered the subunit threshold while keeping all other parameters fixed. As shown in Figure 4E, reducing the threshold transformed the cell’s LSTA profile: originally, the LSTAs tracked the leading edge of the moving bar across time (left), but after lowering the threshold, they became spatially fixed, responding to contrast modulations across the receptive field regardless of the bar’s position (right). This manipulation effectively converted the model from an ON FL–like behavior to an ON S–like behavior, mimicking the difference between high-frequency and low-frequency ON RGC types (see Supplementary Figure 4 for a detailed explanation). Similar manipulations of the other parameters did not produce this switch; while they affected the response amplitude, they did not alter the LSTA dynamics (data not shown).

These results identify the subunit rectification threshold as a key parameter governing whether a modeled RGC is selective to contrast modulations at specific parts of a moving object (i.e., its edges), or responds across the entire receptive field.

### Different ganglion cell types convey parallel and complementary representations of a moving texture

We then asked how RGC types with distinct selectivity profiles contribute to the encoding of moving texture at the population level. We designed a structured moving texture in which a 450 μm-wide bar was divided into three spatial segments—leading edge, middle, and trailing edge—each assigned an independent contrast pattern. Each of the three spatial segments could have two contrast values, leading to 8 different 1D textured bars.These textured bars were shown both to ex-vivo retinae and to the models presented in the previous section.

Then, following previous approaches^8,17^, we used the response of a single cell to simulate a population of identical neurons with different receptive field locations. We composed the 2D texture by vertically stacking the 8 different 1D textured bars, and reconstructed the neural image in response to it (Figure 5A, see Methods). We applied this method both to recorded responses and model predicted responses. This revealed that RGCs with different LSTA behaviors produced population activity patterns that selectively represented different segments of the texture: ON FL population responses enhanced the contrast at the leading edge segment of the texture, both in the model and in the data (Figure 5B), and OFF FT cells responses enhanced the trailing edge segment (Figure 5D). For ON S RGC, it confirmed that these cells are not selective to a particular part of the bar, but rather respond to any contrast change inside their receptive field. Modeling results seem to indicate a weak selectivity of these cells for the central segment of the texture (Figure 5C, left), while in the data ON S responses are more diffuse (Figure 5C, left), showing no particular selectivity for any texture segment.

**Figure 5.**
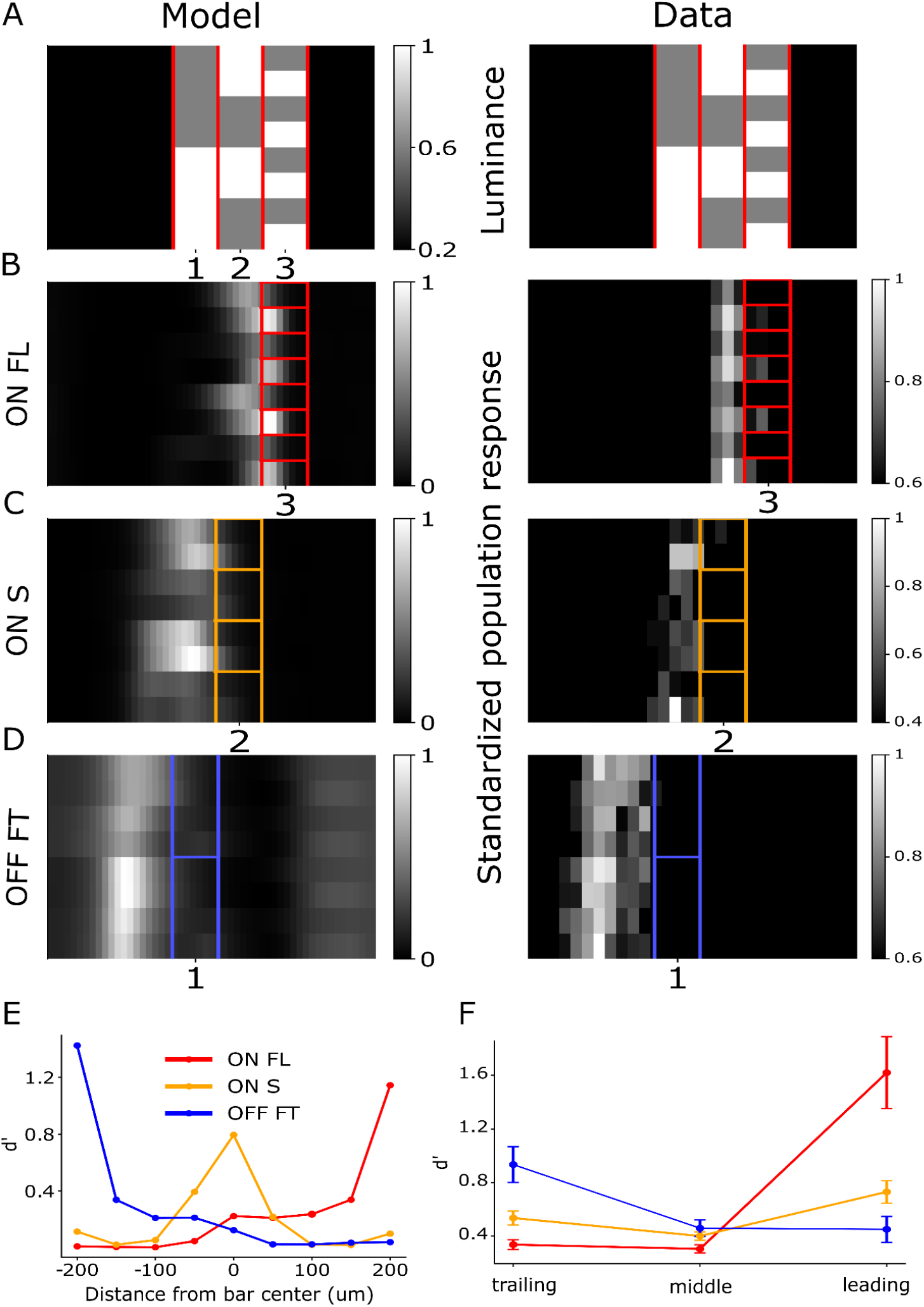
Different ganglion cell types process different parts of a moving texture in parallel. **A.** One frame of the three-part moving texture used to probe how different regions of a moving object are encoded. The 450 μm-wide bar is divided into three segments—trailing, middle, and leading—each assigned an independent contrast pattern. The texture is constructed by vertically stacking eight different 1D textured bars, allowing responses to the 2D stimulus to be inferred from measured 1D responses. The stimulus is repeated twice to ease comparison in the rest of the Figure. **B–D**. Simulated neural images for representative RGCs (model, left; data, right), one from each of the LSTA behavior classes: ON following leading edge (ON FL), ON static (ON S), and OFF following trailing edge (OFF FT). Each panel shows the standardized response of a population of identical copies of a given cell with shifted receptive fields^17^, at a specific time across the texture sweep. Colored boxes indicate the spatial location of the corresponding texture segment (red: leading edge, orange: middle, blue: trailing edge). **B**.ON FL cell population activity tracks contrast in the leading segment of the texture. **C**. ON S population activity reflects a more diffuse pattern, partially matching both the center and leading segments. **D**. OFF FT population activity follows contrast in the trailing segment. **E**. Discriminability index for a moving texture with 9 segments, computed from model predicted responses. The index measures how much a cell’s response varies with changes in contrast at a specific texture segment. ON FL cells show peak discriminability for the leading edge, ON S for the center, and OFF FT for the trailing edge. Similar to panel F but with 9 distinct segments of 50 µm instead of 3 segments of 150 µm. **F**. Average discriminability of the three texture segments across recorded RGCs of each LSTA type. Each data point represents the average d’ for a specific texture region and RGC group. While the ON FL and OFF FT groups show high selectivity for the leading and trailing edges respectively—consistent with modeling results in panel E—the ON S group shows more uniform, moderate discriminability across all three regions.

To quantify these effects, we computed for each cell and texture segment a discriminability index (d’, see Methods). This index quantifies for a given segment how much the response of a cell to the texture varies in function of the contrast content of the segment. The higher the index, the more the cell responses change when only the contrast of the selected texture segment changes. ON FL cells showed peak discriminability for the leading segment, OFF FT for the trailing segment, and ON S cells exhibited more uniform but lower discriminability across the texture, consistent with their static LSTAs (Figure 5F).

Because the three-segment texture is spatially coarse (150 μm per segment), we leveraged the model to probe finer spatial selectivity. We repeated the discriminability analysis using model responses to a nine-segment texture of equal total width but 50 μm segments. This increased resolution revealed that different types of RGCs are selective to non-overlapping segments of the moving texture on a spatial scale (50 µm) that is 3-4 times smaller than the typical RGC receptive field size.

These results suggest that distinct RGC types encode complementary, spatially localized information about moving textures in parallel.

## Discussion

Thanks to the center-surround organization of their receptive fields, ganglion cells are able to enhance edge contrast in a static visual scene, a first step towards image segmentation. For moving objects, it has been shown that this center-surround organization can also help for motion detection. However, the retina is more than an array of linear filters with a mexican hat profile ^23^. In particular, moving objects tap into several non-linear mechanisms in the retina ^19^. It was thus unclear how these non-linearities would impact the encoding of moving object boundaries and therefore the segmentation of different moving objects.

Our results show that specific types of ganglion cells dynamically tune their spatial selectivity to enhance contrast near the edges of a moving object. When measuring LSTAs with our perturbative approach, we found that some tracked the texture content near the leading edge of the object, while some tracked the one near the trailing edge, and others remained static. This behaviour was due to the strength of the non-linearity that rectifies the subunit output.

This change in spatial selectivity differs between cell types, and has a consequence at the population level. Some cell types enhance contrast changes near the leading edge, others near the trailing edge, and others have a constant spatial selectivity no matter where the bar is.

While it is clear that motion segmentation is not solved by the retina, our results show that these cell types provide parallel and complementary representations of a moving object to the brain that can be used as primitives for motion segmentation.

### Coding the content of moving objects

Previous works have studied how moving objects are encoded by retinal ganglion cells. However, they focused on complementary questions. The encoding of motion direction has been addressed by many studies of direction selective cells^7^. Several works have studied how the position of a moving bar is encoded, during steady motion^8,9,24^, motion reversal^25^, motion onset^19^, or random motion ^18,26^. However, they did not study how the content of the object, i.e. the texture, is encoded by ganglion cells. Most of the studies who focused on texture encoding were based on static stimuli^21,5,27,14^, with motion cues often studied separately. This is also the case beyond the retina, where texture and motion cues are often assumed to be processed separately ^28^. Here we show that moving textures uncover specific encoding strategies in specific types of ganglion cells that could not be found using either uniform objects or static textures.

Our modeling study suggests that a model needs to combine both subunits and gain control to predict the responses to moving bars and the LSTA dynamics. Although we could have retrained the original LNLN model directly on the bar stimulus instead of adding a gain control, doing so would have compromised its performance on the white noise data, undermining its ability to generalize across stimulus classes. By contrast, extending the model with a gain control stage introduced only one additional free parameter and allowed the model to generalize to moving stimuli while keeping the ability to predict responses to white noise. Having these two components thus allowed having a model that can generalize and predict the responses to different stimulus statistics: white noise, moving bars, and perturbed moving bars.

We aimed at finding the simplest model that could reproduce our experimental results to make the interpretation of the model parameters as simple as possible. This was possible with a gain control only at the level of ganglion cells. However, the retina has been shown to adapt more locally than at the ganglion cell level, e.g. at the photoreceptor level^29^ and possibly at the bipolar cell level. This evidence calls for models that comprise more local gain controls^20^, but given the simplicity of the moving bars used here, including them in the model architecture proved to be unnecessary (data not shown).

### Motion anticipation

Our results address an orthogonal and complementary question to the previous studies of motion anticipation in the retina. When responding to a steadily moving object, the retina seems to have a mechanism for partially compensating the synaptic transmission delays that limit the animal’s reaction time to motion onsets ^8^. This compensatory mechanism was shown by highlighting that the neural image of some OFF RGCs populations is aligned with the leading edge of a moving bar instead of lagging behind it. This was true for fast OFF and other OFF RGCs in the salamander retina, and brisk-transient OFF, brisk-sustained OFF and local edge detectors in the rabbit retina. In the mouse retina, the same effect was observed ^10^.

Our results suggest that different types may have different levels of anticipation. In figure 5, the position of the neural image peak relative to the texture was different for each RGC type. For ON FL ganglion cells, the peak was aligned with the leading edge of the texture, while for OFF alpha transient (OFF FT) it seemed to lag behind the bar’s trailing edge. This suggests that the different types might not only represent contrast variations on different parts of a moving texture, but also do so with different temporal lags. We observed that the RGC types that were shown to detect the leading edge seemed also the ones with the strongest motion anticipation, while the other types showed neural images lagging behind their preferred texture segment, but confirming this trend would require testing it for different speeds.

### Mechanisms to process moving textures

Our results show that different cell types have different LSTA dynamics. Presumably, this reflects differences in the circuits of each type.

Our modeling study gives insight on the putative mechanisms that allow the retina to track contrast changes at the edges of a moving object. The model architecture mirrors the structure of several known components of the retinal circuit. The subunit corresponds to the processing performed by bipolar cells. The filter of the second stage reflects the spatial integration performed by ganglion cells. The final rectification threshold mirrors the spiking threshold of ganglion cells. The gain control likely reflects the combined effect of several adaptation mechanisms (see Discussion in ^19^).

The rectification of the subunit may be implemented by the synapse from bipolar to ganglion cells. Several works have suggested that this synapse implements a rectification that makes spatial integration non-linear ^30,31^. However, other mechanisms could implement this rectification, e.g. non-linearities in the compartment of ganglion cell dendrites^32^. Our modeling study predicts that decreasing the rectification threshold could remove the tracking behaviour of LSTAs. Amacrine cell inhibition of the bipolar cell terminal could also modulate the rectification threshold. Depolarizing the bipolar cell could result in the same effect. Both manipulations would require to know which types of bipolar cell are inputs to the ganglion cells showing tracking behaviour, or to know the amacrine cell types that inhibit the terminal of these bipolar cells. Future studies will thus have to elucidate the connectivity of this circuit, and to implement methods to modulate the activity of specific cell types in the retina^33^. In summary, our model-based insights point to a testable hypothesis: differences in edge-tracking versus static selectivity across RGC types may arise from cell-type–specific tuning of subunit thresholds. By tying functional response patterns to a specific and biophysically plausible nonlinearity, our findings provide a mechanistic link between retinal circuitry properties and the encoding of motion-related texture features.

The LSTA dynamics could not be simply related to the transient-sustained nature of the cell type. One hypothesis could have been that transient cells would only respond when a moving object enters their RF, generating LSTAs only around the leading edge, while sustained cells would respond during the whole time the bar overlaps with the receptive field, producing more static LSTAs. However, our data does not support this hypothesis. For example, ON transient cells showed clear transient responses to full-field flashes, but did not track the leading edge significantly (Figure 2C). Conversely, ON and OFF alpha sustained cells, which had sustained responses to step stimuli, showed edge-tracking behavior (Supplementary Figures 2 and 3). These findings suggest that spatiotemporal selectivity during motion is not solely a consequence of the transient/sustained nature of the cell, but likely arises from dedicated circuit mechanisms that modulate sensitivity depending on the object’s motion, where the non-linear rectification at an intermediate stage of the retinal circuit plays a key role.

### A first step towards segmenting moving objects

Rodents are able to modify their behaviour in the presence of visual objects. They can perform object recognition^34,35^ and use it for spatial navigation^36^, or catching prey^37^, although their ability to segment visual objects might be more limited than for primates, and may require contrast differences between figure and background ^38^.

Segmenting visual objects is thus a challenging task^39^, but useful for many behaviours. This requires assigning an object identity to all the pixels that constitute a surface, a feat that is not achieved in the retina. However, the retina can perform a pre-processing step to extract computational primitives for segmentation, i.e. features relevant for this task. In this sense, the retina contributes to the segmentation of a static image, by enhancing the edges of visual objects. This is done thanks to the center-surround structure of the receptive field of most ganglion cells.

Here we show that specific types of ganglion cells can selectively enhance contrast changes near moving edges, and be insensitive to contrast changes in other parts of a moving object. This property could not be derived solely from the receptive field center-surround structure ^12^ as a specific non-linear subunit rectification was necessary. Our work shows that critical non-linearities give a specific spatial selectivity that allows tracking of moving textured edges, a key step for solving discontinuity and beginning to segment moving objects.

To illustrate how these types of cells could be useful in a more natural context, we designed a toy model that includes all the key components of the model we have used above, including the subunit rectification and the gain control, but integrating over a 2D space. We simulated the response to a drifting natural image (Figure 6A) of an array of cells that would be similar to a ON leading edge cell like the ones we characterized above (Figure 6B and supplementary video V2). The population response clearly enhances the boundary of the moving object (the wing of the butterfly, Figure 6B). In comparison, a linear model was clearly less specific (Figure 6C). The activity in response to the edge of the wing also represents the variation of contrast around the edge, with a decrease corresponding to the dark regions.

**Figure 6.**
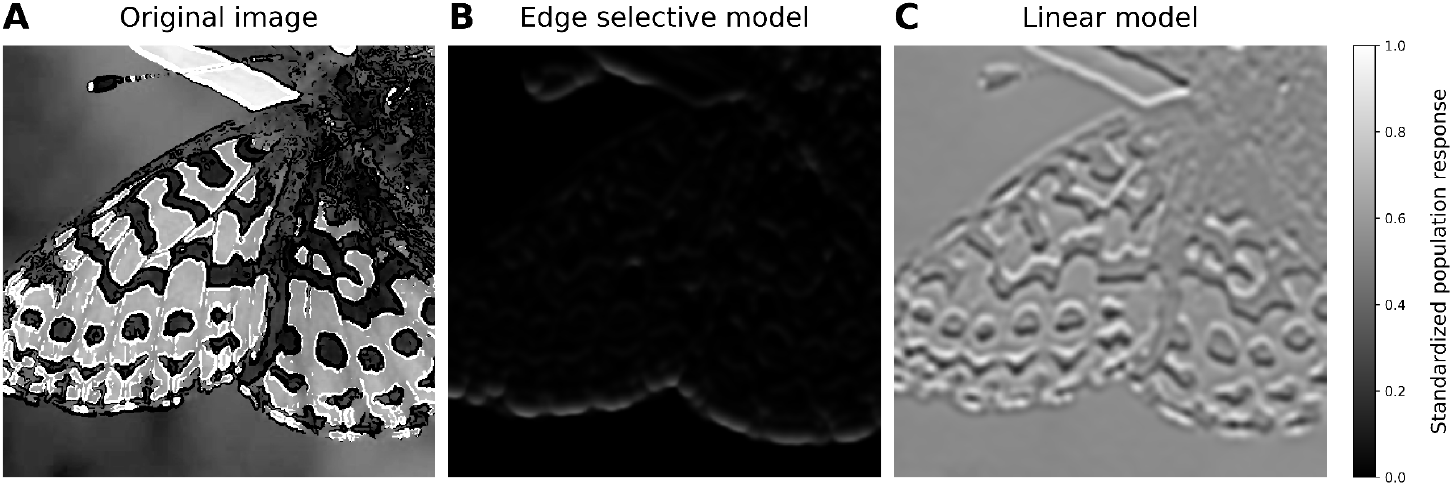
Moving texture edge extraction in a naturalistic stimulation context. **A.** Example frame from a downward-drifting butterfly stimulus. See Supplementary Video V2 for the full sequence. **B**. Output of the leading-edge selective model. The stimulus is first convolved with a spatiotemporal difference-of-Gaussians filter representing subunit receptive fields, followed by rectification. The rectified signal is then convolved with a larger Gaussian filter to simulate the population response of a regular grid of ganglion cells and subsequently modulated by divisive gain control (see Methods). Responses are shown after standardization to the range [0,1]. **C**. Output of the corresponding linear model, in which the rectification and gain-control stages are omitted while all other filtering stages are identical. Responses are standardized independently. All the parameters of the models were chosen by hand in arbitrary units.

The output of this model does not allow to segment objects per se, a task that is performed beyond the retina: for example, it does not distinguish between edges inside an object and border between two objects, and it does not code for border ownership. However, it will make it easier to track boundaries across time, since they are clearly enhanced in this representation.

By representing specifically the texture near the moving edge, specific types of the retina make the tracking of object boundaries across time easier, because the boundaries can be matched across time and space more easily once they have been isolated by these types. This representation could thus be useful for motion segmentation performed in downstream areas. Identifying the downstream areas where they project could also help narrowing down for which visual tasks they can be useful.

Several studies suggest that cortical visual areas, analogous to the ventral stream in primates, represent visual objects with some degree of invariance to position and rotation ^34^. Objects also modulate the tuning of neurons in areas related to spatial navigation ^36^. However, in most cases, texture and motion processing have been studied independently, and are often assumed to be represented by different areas of the brain ^40,41,28^. To test for a causal role of the representations we characterized in this study, it would be necessary to inactivate the cell types that perform this edge enhancement and test if this impacts the capability of the animal to segment moving objects.

More generally, the segmentation problem is often studied in cases where only one type of cue is available - e.g. motion, contrast, or texture. Our results uncover specific non-linear computations where spatial selectivity is dynamically shaped by motion. This suggests that the visual system may have specific non-linear computations to process combinations of these different cues, that would be missed if they are studied in isolation.

## Acknowledgements

We would like to thank Fred Rieke for critical comments on the manuscript and Matias A. Goldin, Ulisse Ferrari and all the members of the Marre lab for helpful discussions. O.M. was supported by the ERC Consolidator grant DEEPRETINA (101045253), ANR grant Chaire Industrielle MyopiaMaster ANR-22-CHIN-0006, ANR grant ANR-18-CE37-0011–DECORE, ANR grant ANR-20-CE37-0018-04–Shooting Star, ANR grant ANR-22-CE37-0033 NUTRIACT, ANR grant project ANR-22-CE37-0016-01 PerBaCo. ANR grant project RetNet4EC (O.M., M.C.), AVIESAN-UNADEV, Retina France and Programme Investissements d’Avenir IHUFOReSIGHT 497 (ANR-18-IAHU-01). M. C. was supported by ANR project ANR-22-CE92-0015. M. C. thanks Qube Research & Technologies (QRT) for their financial support via the project DISSENSATION. T. B. was supported by the PhD programme of the Fondation pour la Recherche Medicale (FRM). S.V. was supported by the European Union’s Horizon 2020 research and innovation programme under the Marie Skłodowska-Curie grant agreement No 861423. O.M.’s lab is part of the DIM C-BRAINS, funded by the Conseil Régional d’Ile-de-France.

## Author contributions

Conceptualization, SV,TB,MC,OM; Data acquisition, TB; Data curation, SV,TB; Formal analysis, SV; Funding acquisition, OM,MC; Methodology, SV,TB,MC,OM; Supervision, OM; Writing – original draft, SV,TB,MC,OM; Writing – review & editing, SV,MC,OM;

## Declaration of interests

The authors declare no competing interests.

## Methods

### Recordings

Recordings were performed on C57BL6/J adult mice of either sex. The animals were housed in enriched cages with ad libitum food and watering. The ambient temperature was between 22 and 25 °C, the humidity was between 50 and 70% and the light cycle was 12–14h of light, 10–12h of darkness. The experiments were performed in accordance with institutional animal care standards of Sorbonne Université. After the sacrifice of the animal, the eye was enucleated and rapidly transferred to oxygenated Ames medium (Merck, A1420). The retina was extracted from the eye cup and the vitreous was carefully removed. One quarter of the retina was lowered with the ganglion cell side against a multi-electrode array with electrodes spaced by 30 µm, as previously described ^42^. During the recordings, the Ames’ medium temperature was maintained at 35-37 °C. The raw voltage traces were digitized and stored for offline analysis using a 252-channel preamplifier (MultiChannel Systems, Germany) at a sampling frequency of 20kHz. The spikes of individual ganglion cells were isolated using Spyking Circus^43^. Subsequent data analysis was done with custom-made Python codes.

### Visual stimulation

Visual stimuli were presented using a white LED (MCWHLP1, Thorlabs Inc.) and a Digital Mirror Device (DLP9500, Texas Instruments) and focused on the photoreceptors using standard optics and an inverted microscope (Nikon). The mean light levels used for all stimuli are in the range of photopic vision.

### Cell type characterization stimuli

We displayed a set of standard visual stimuli to characterize RGC types. First, to measure the classical RF through reverse correlation, we presented a two-dimensional random white noise stimulus at 30 Hz. The check size was 105 µm and the stimulus duration was 50 minutes. Then we presented the full field chirp stimulus introduced in ^13^. It was played at 50 Hz, containing 20 repetitions of 32 s length. Finally we presented drifting gratings moving in 8 directions. The gratings had a speed of 479.5 µm/s, at a spatial period of 959 µm and at 50% Michelson contrast (0.75-0.25 luminance). The 8 directions were repeated 4 times and randomly interleaved. Each repetition lasted 10 s, preceded by 2 s of gray, the temporal period being 2 s. The responses to these three stimuli were used to cluster ganglion cells in the functional types introduced in ^13^. Details of the clustering are reported in ^14^. To estimate the cells’ classical 1D STA, we presented a 1-dimensional white noise stimulus at 40 HZ for 50 minutes. The size of the stripes was 31.5µm. A two-dimensional STA (x and time) was sampled using 21 time samples.

### Perturbed moving bar

The stimulus was composed of a moving bar 451.5 µm wide, moving at the constant speed of 840µm/s superimposed to dim 1D white noise patterns 31.5 µm in width. The perturbations had an amplitude of 30% (0.2 luminance on top of a 0.6 bar on a 0.2 background). A single bar sweep lasted 69 time frames at 40 Hz (1.725 s) and was repeated 2050 times. The random perturbations changed at every time point across all the repetitions. The amplitude of the white noise patterns was chosen to elicit a small but visible change on the average ganglion cell response compared to the response to the unperturbed moving bar (see calibration in Supplementary Figure 1). The bar was shown both moving from left to right and from right to left. To avoid adaptation to one direction of motion, we interleaved randomly the two opposite directions of motion across repetitions.

### Calibration of perturbation amplitude

We wanted to find the minimal perturbation amplitude that produced a discernible spike count difference between the response to the uniform moving bar and to the corresponding perturbed moving bar. In a calibration experiment, we presented a uniform moving bar superimposed with a fixed perturbation that could assume 12 different amplitudes. We repeated each perturbation sequence 140 times, presented in a randomized manner. We chose a perturbation amplitude of 30%, where 100% corresponds to the contrast of the bar compared to the background illumination. This value produced an average change in firing rate of 30 Hz across the ganglion cells psths measured over the 1.725 s of the bar sweep (Supplementary Fig. 1).

### Textured moving bar

The stimulus was composed of a 451.5 µm bright bar moving in either direction at a speed of 840 µm /s. The bar was presented either uniform, or with a contrast increase covering either its leading third, middle or trailing third. The contrast increase had an amplitude of 30% (0.2 luminance on top of a 0.6 bar on a 0.2 background). All conditions were randomly interleaved.

### Subunit model

For modeling the 1D white noise we used a single-cell CNN model composed of 1 convolutional layer followed by a readout. Both the convolutional kernels and the readout are spatiotemporal (2D, 1 dimension of space and 1 of time). The model takes as input a snippet of the spatiotemporal stimulus *S*(*t*) at time *t*. The snippet spans the whole stimulus space in x (96 pixels) and a time lag τ_*_ of 25 time points from *t* to *t* – τ_*_. This snippet *S* _*xτ*_ is convolved in parallel with two convolutional kernels *K*_*c*_ with *c* = {*ON, OFF*}. Then the outputs of these kernels are rectified with a non-linearity ϕ_*c*_ (different per each channel) and they are projected onto a common dense layer *v*_*xτ*_. This produces a set of *c* = 2 numbers that can be averaged together with some weights *w*_*c*_ into one number. Finally, this one number is passed again through a non-linearity ψ, to produce the predicted response:

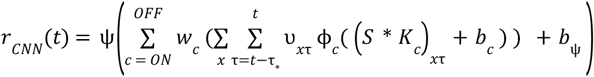

where *x* and τ index space and time lag, respectively, and *c* indexes the convolutional channels. We choose ϕ_*c*_ (*x*) to be a relu and ψ(*x*) to be an elu, common non-linearities in machine learning:

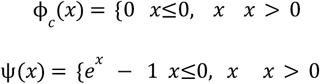

Because of the last non-linearity, the predicted response of the model is given as *r* + 1. During training, the convolutional kernels are initialized with bipolar cells receptive field data (^22^, figure 2G) and kept frozen, while the three nonlinearity biases *b* _*ON*_, *b* _*OFF*_, *b* _ψ_ and the readout weights *v* _*xτ*_ are trained through gradient descent (Adam optimizer). While in ^22^ authors identified 14 distinct bipolar cell types, we simplified the subunit layer by averaging receptive fields separately for ON and OFF bipolar cells, resulting in a single ON subunit and a single OFF subunit. Fixing these subunits conferred several advantages. It grounded the model in biological measurements, ensuring mechanistic interpretability by directly linking model components to specific retinal interneurons. It also greatly reduced the number of free parameters, simplifying training and improving model stability. Because the subunits were the same across all cells, model comparisons across cell types were more meaningful, with differences in behavior arising from downstream integration and rectification rather than subunit structure. The training is done through TensorFlow custom code to minimize a regularized negative poisson loss of the form:

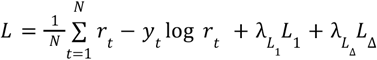

evaluated on a validation set that was held out during training (20% of the un-repeated database). In the previous equation, *y*_*t*_ is the recorded response at time *t, r*_*t*_ is the predicted response by the model at time *t*, and 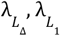 are the hyperparameters which control the importance of the Laplacian *L*_Δ_ and *L*_1_ regularization terms applied to the readout with 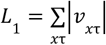 and 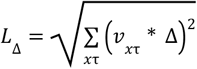where 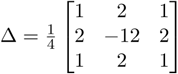

The modelled cells were 79 across 4 retinas and were chosen among the ones that presented visible LSTAs on the perturbed moving bar and were stable across the 1D white noise stimulation. We could assess stability because we interleaved random 1D white noise sequences with repeated sequences. Given n responses to the same repeated sequence, {*y*_1_, …, *y*_*n*_}, we averaged them over odd and even numbered trials to get two estimates of the mean response, 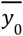 and 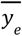. We defined the reliability as the pearson correlation between these estimates, 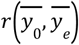. This index goes from -1 to 1 and values close to 1 mean that a cell is spiking consistently. We considered only cells with reliability greater than 0.8. These repeated sequences were also held out during training and were used as a testing set. After training, the performance of the model was evaluated by calculating a noise corrected pearson correlation between the model prediction 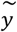 and the data of the form:

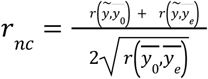

We selected for further analysis only the 56 cells that had a performance above 0.5 (Figure 3C).

With this subunit model trained we tried to predict the cells’ responses to a uniform moving bar of the kind used for the LSTAs estimation but without perturbations. In our dataset we had 30 repetitions of a uniform bar, but for robustness we decided to complement these data with the PSTH of the randomly perturbed moving bar averaged across the 2050 repetitions. Since the perturbations were white and random, their average gave a similar but much less noisy estimate of the response of the cell to the unperturbed bar. The subunit model performed well for many cells but for others it predicted the wrong response onset and response duration due to the time correlations present in the continuously moving bar stimulus. To fix this, instead of retraining the whole model, we decided to add to the scaffold trained on white noise a gain control mechanism for time adaptation^19^. In this new model the predicted response at time *t*:

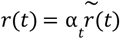

Where 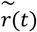 is the response predicted by the previous CNN model and

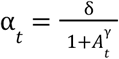

with

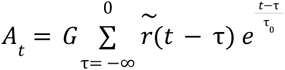

This gain control layer in principle introduces 4 new free parameters in our model *G*, τ_0_, *γ*, δ. Ideally, those should be fitted from the data in the same way we fit the other parameters. In practice, since they are part of a feedback mechanism, their training is unstable through gradient descent. For this reason, we decided to pseudo-train *G* by grid-searching the values that for each cell increased the value of the noise-corrected Pearson correlation between the model predictions and the moving bar data. τ_0_ and *γ* were found to be underconstrained with respect to *G* and so their value was fixed at 400 ms and 1s respectively. Finally, the scaling factor δ was necessary because since the optimization of *G* was performed on a correlation measure, sometimes it caused the amplitude of the predicted response to be very different from the one observed in the data. So after training *G*, the psth amplitude was rescaled by δ to minimize the mean square error between the data *y*(*t*) and the model predictions *r*(*t*):

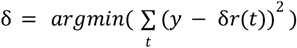

The solution can be analytically found to be:

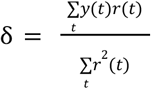

In the end, the time gain control only added one free parameter, the strength *G*.

### Position-dependent spike triggered average computation

In order to measure LSTAs from the data, we reverse-correlated the cell responses to the perturbed moving bar. The bar had uniform velocity v and a single bar sweep lasted 1.725 s, which at 40 Hz results in 69 time frames. Let us index those frames with the letter t. Since the bar is 1D, the sweeping bar movie can be represented with a 2D array *S*_0_ of dimension (96,69) where 96 is the number of pixels in space and 69 the number of time points (like in Figure 2A, left). Let us define 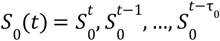 where τ_0_ = 12 and 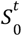 is the column of *S*_0_ corresponding to time t. In each column 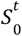, the moving bar has a different position and a different noise superimposed on it. The bar sweep was repeated 2050 times, each time superimposed with a different random perturbation. Let us index the repetitions with the index *r*. In this way, we can write the whole perturbed moving bar stimulus as *S*(*t, r*) = *S*_0_(*t*) + Δ(*t, r*) with *t* ∈ {τ_0_, …, 69} where Δ(*t, r*) indicates the perturbations. The ganglion cell responses during each bar sweep were binned with 25 ms bins, resulting in a response of 69 bins, one per bar position. We can therefore write the response of a single ganglion cell as *y*(*t, r*) without ambiguity. This vector has dimension (69,2050). Note that, given the position of the bar at time *t*_0_, *S* (*t*_0_, *r*) is a set of 2050 bars in the same position that differ only for a dim random perturbation, and therefore it provokes a set of 2050 responses *y*(*t*_0_, *r*) that are similar to each other but not identical. We can therefore reverse correlate this set of responses to obtain a receptive field estimation: : 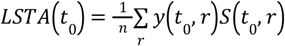 where *n* = 2050. *LSTA*(*t*_0_) is a 2D array of shape (96, τ _0_) where 96 indexes space pixels and τ_0_ is the time points of lag from *t*_0_. Then per each cell, we measured LSTAs for different times *t* (examples in Figure 1G). Having chosen a τ_0_ = 12, we had 57 2D filters for each cell. To compress this representation and effectively convey the spatiotemporal LSTA behavior of a cell at a glance, we decided to fix per each cell a certain τ and to stack the 1D slices of each of the 57 LSTAs at time lag τ together to obtain a 2D summary of shape (57,96). Examples of this plot are reported in Figure 1H and 2B). The τ was chosen automatically as the time lag at which the time course of the LSTA in absolute value was greatest.

The predicted LSTAs of the model could be obtained by differentiating the trained model with respect to the stimulus and by calculating the model gradient around the different bar positions at different time points:

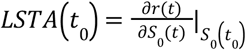

### LSTA denoising model

Taking the reverse correlation of a dim white perturbation on top of a much more salient moving object gives rise to a detectable (provided many repetitions) but very noisy signal. For this reason, in order to do quantifications on the LSTAs, it was necessary to denoise them. Previous attempts to tackle similar problems successfully used Gaussian processes to this end^44^. We therefore used a Gaussian process similar to the linear one described in ^45^, but extended to spatiotemporal stimuli. In order to do that, we modified the Bayesian prior as suggested in ^46^.

This led to a model where the spike count *r* is assumed to be drawn from a Poisson distribution with mean:

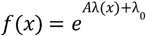

where x is an *n*_*x*_ -d stimulus vector, A and λ_0_ are scalar gain and bias terms, and λ(*x*) is a weighted sum of *J* units:

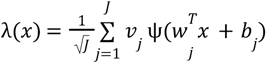

where *w*_*j*_ are *n*_*x*_ -d vectors of linear weights, *b*_*j*_ is a scalar bias term, *v*_*j*_ is a scalar weight and ψ is a threshold non-linearity (relu). Making some a priori assumptions on the distribution of the parameters *w*_*j*_, *b*_*j*_ and *v*_*j*_, and considering the limit *J*→∞, we can use the central limit theorem to write λ(*x*) as a Gaussian process, with a closed form for the mean and covariance, *K*(*x, x*’) that only depends on a handful of parameters that parametrize the prior on the weights *w*_*j*_. In this way, we impose a strong regularization to our model. We can then fit those parameters to maximize the posterior probability of *w* given the white noise perturbations as stimuli x and the perturbed bar responses as *f*(*x*) (details in ^45^). Once fitted, the model can be differentiated to obtain a gradient *g* that is a denoised version of the LSTA.

We fitted a separate model for each one of the tenths of LSTAs that we measured per each recorded cell. Since λ(*x*) is a gaussian, *g*, its gradient, is also a gaussian *g*∼*N*(μ_*g*_, Σ_*g*_). Σ_*g*_ in particular estimates the pixel-wise uncertainty of the model in its gradient prediction. This allowed us to define a quality selection criterion for our denoised LSTAs. We wanted to know whether our predicted LSTA was significantly different from 0 and so it should be trusted, or discarded. If we assume the LSTA pixels to be distributed with a gaussian, we can compute the quantity 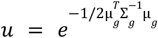 that estimates the probability that the null hypothesis LSTA = 0 is true. We therefore kept only the LSTAs whose uncertainty *u* was lower than 10 ^−9^.

### Neural Images

The neural image method^17^, is a way of rearranging in space and time the responses of a single neuron to effectively simulate the response of a population of identical neurons that tiled the visual field homogeneously. This was possible in our analysis because in the case of a continuously moving bar space x and time t are correlated. We could therefore simulate a grid of cells in different places along x by shifting the psth of a cell with RF center in *x*_0_ in time of an amount 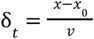 where v is the velocity of the moving bar. Also, since our stimuli moved in a straight line along x, for the y-axis the RF of the recorded cells was assumed to be 1 pixel wide. With this assumption, the 2D moving texture response could be obtained as a stack along y of responses to 1D moving textures.

### Tracking index

We wanted to have an index to quantify whether and how much the selectivity of our cells was affected by the presence of a bar edge. Because of our way of measuring selectivity though LSTAs, this resulted in determining whether the positions of the LSTAs measured for a cell over time were correlated with the position over time of the bar edges. So we decided to adopt as a tracking index the fraction of variance in the LSTA position over time that could be explained by the bar. Within a chosen window of time {*t*_1_, *t*_2_, …, *t*_*n*_} this measure writes:

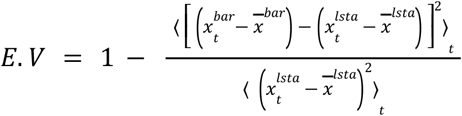

where 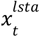 and 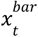 are the positions of the LSTA and the bar edge at time t and 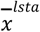 and 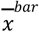 are the average positions of the LSTA and the bar edge across the chosen time window. This index will have value 1 for the cells whose LSTAs faithfully follow the bar edge and will have a value around 0 for cells whose LSTAs are static in time. The index was calculated for every possible window of time larger than 150 ms across the bar sweep and the window with the highest index value was chosen. This allowed in the case of the edge-tracking cells to automatically detect the window of time in which the bar was entering or exiting their receptive field. For the cells with static LSTAs the choice of window had little importance. Note that since this is a correlation measure and the correlation between the two bar edges positions in time is 1, the tracking index will have the same value for the leading and the lagging edge.

Then we wanted to determine whether the cells of one specific type were all significantly tracking the bar edges or not. To do this, we calculated for each cell, besides the measured tracking index, 1000 other null tracking indexes obtained by randomly shuffling the LSTAs in time. We then constructed for each RGC type a true and a null tracking index distribution by pulling together the true and the null tracking indexes of all the cells of the type. Finally, we compared the two distributions with a two-sample Kolmogorov-Smirnov test and we deemed as significantly edge-tracking only the RGC types whose p-value was smaller than 10 ^−4^.

### Discriminability index

The aim of this index is to quantify for each texture segment whether its contrast modulation is affecting the firing rate of a given recorded cell. Since the contrast modulations take only two values, either 1 (maximal luminance of the screen) or 0.6, for each texture segment the cell responses can be divided into responses to high luminance (white,1) *r*_*w*_ (*t*) and responses to lower luminance (gray, 0.6) *r*_*g*_ (*t*). We define our discriminability index as:

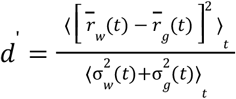

where 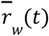 and 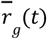 are the averages and *σ*_*w*_(*t*) and *σ*_*g*_(*t*) are the time specific standard deviations across conditions of all the responses to white and grey respectively. Note that since we are voluntarily agnostic to which segment of the texture the responses are caused by, when we divide them into white and grey responses for a given segment, we only do so based on whether their stimulus was grey or white in the selected segment. Therefore, for the segments to which the cell is not selective, the division into white and grey is random and so 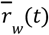 and 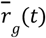 will be almost the same, making the numerator close to zero. Conversely, if the texture segment is meaningful for the cell, then 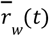 and 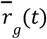 will be different from each other making d’ larger.

## Supplementary Figures

**Figure S1:**
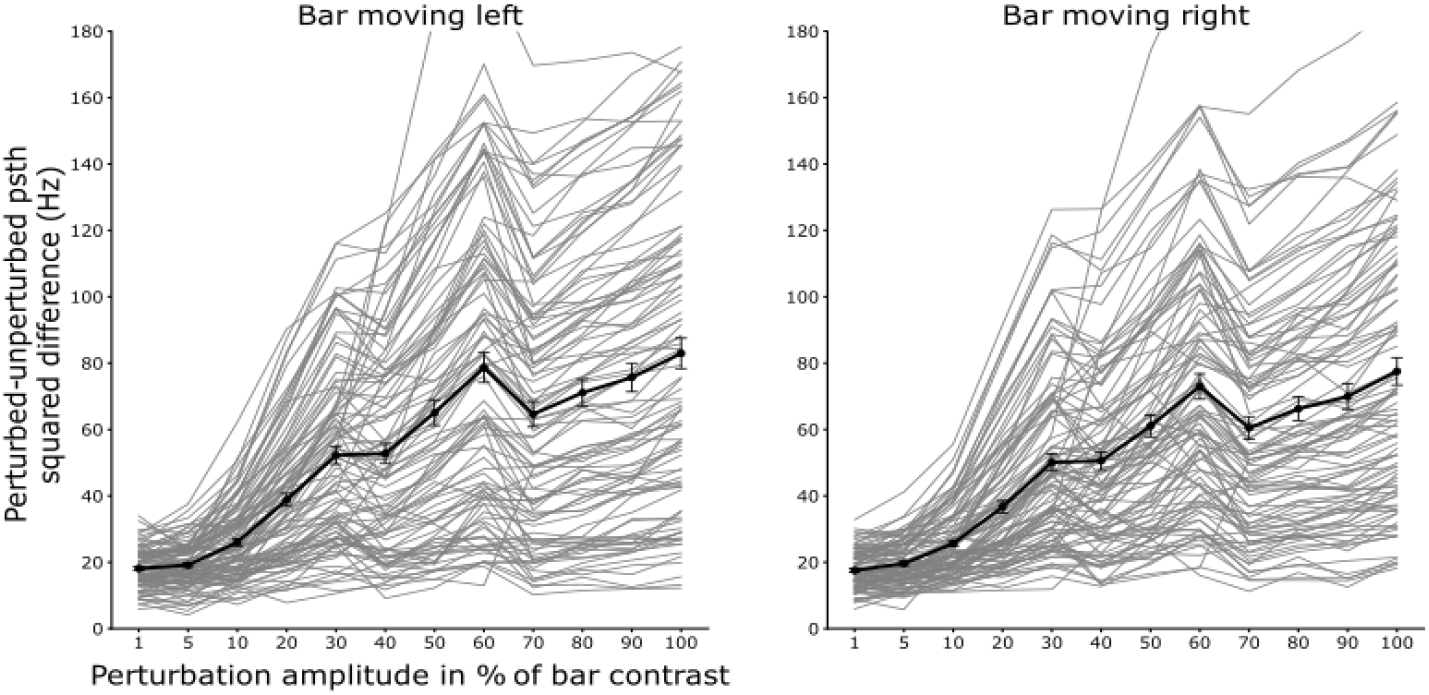
Perturbation contrast calibration. A single perturbation with 12 different contrasts is superimposed on a uniform moving bar either moving left (left panel) or right (right panel). Every grey line is one cell and the values reported are the average mean square difference between the response to the perturbed and the unperturbed bar for the 12 different contrasts. The black line is the average curve across 120 recorded cells. The error bars for every point are the S.E.M across cells. We chose the 30% contrast (30% of the contrast value of the bar).

**Figure S2:**
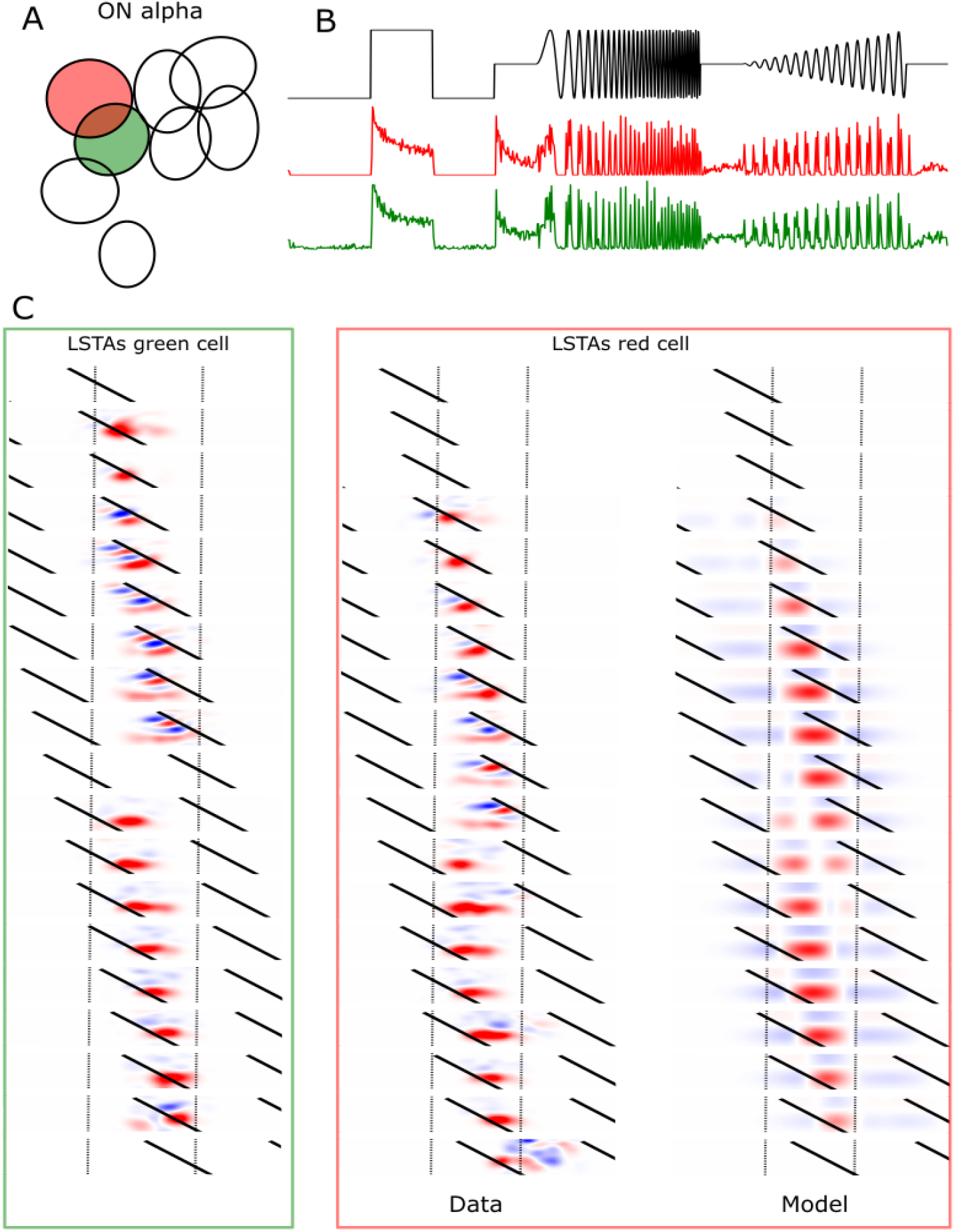
ON alpha RGCs show selectivity to both edges of the moving bar. **A**. Mosaic of ON alpha RGCs. **B**. Chirp responses of two example cells. **C**. Left and middle column: LSTAs modulation behavior of the two example cells in A. Note that cells of the same type show same LSTA behavior. Right column: Pseudo-trained model prediction (see Supplementary figure 4 for model details). The model can qualitatively reproduce the observed behavior for an intermediate subunit threshold value.

**Figure S3:**
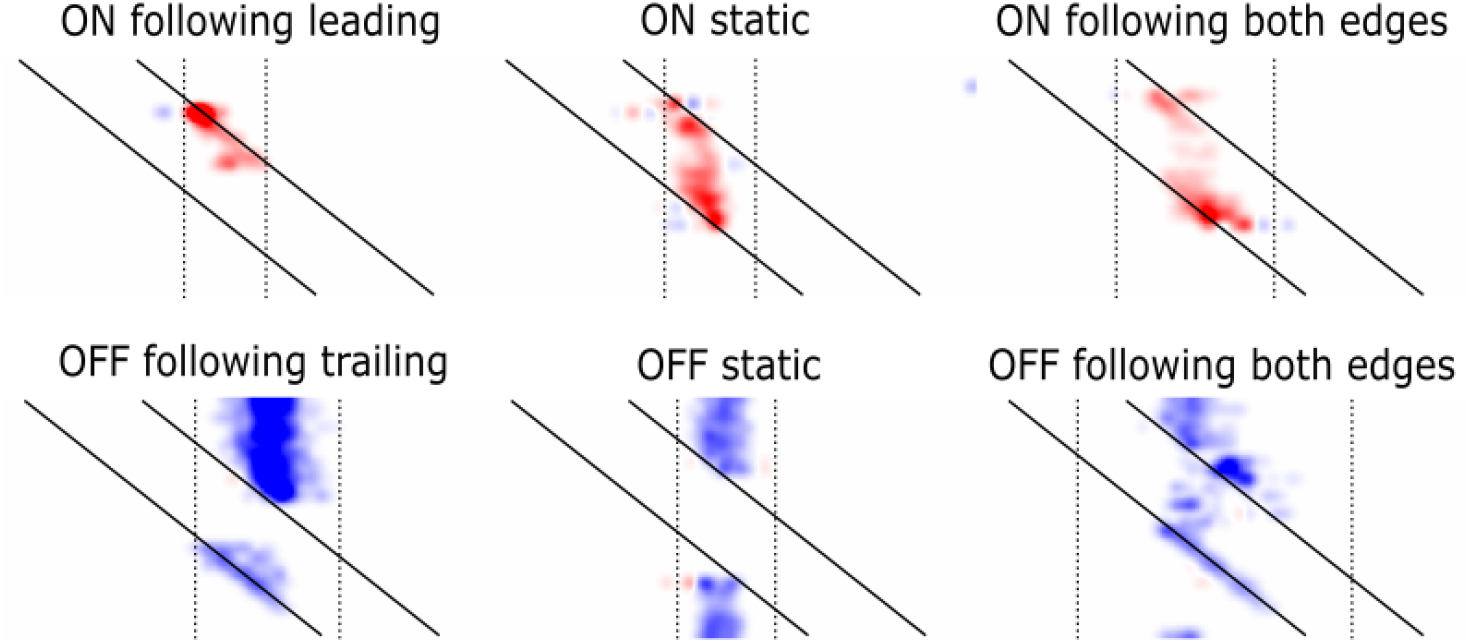
The 6 LSTA behaviors we identified among pure ON and pure OFF cells. Here are reported the LSTAs of 6 example cells, one from each behavior. The example cells reported here are different from the ones in Figure 2 to show more raw data but they are from the same functional types. To know the types associated with each behaviors refer to Figure 2C. ON-OFF cells show a mix of these behaviors (data not shown).

**Table T1:** Are the model parameters significantly different for cells with different LSTA behaviors? In the table are reported the p-values of a Welch’s t-test (t-test that allows different distribution variances for the two samples). The modelled cells were divided into ON FL, ON S and OFF FT according to their chirp responses, not according to their LSTA behavior. The only comparisons for which there was a significant difference were ON FL-ON S and ON S-OFF FT in the case of the subunit threshold. This points to the decisive role of the subunit threshold in determining an edge-tracking behavior (if high) or a static behavior (if low).

|  | ON FL – ON S | ON FL – OFF FT | ON S – OFF FT |
| --- | --- | --- | --- |
| Subunit threshold | 5.7e-4 | 0.63 | 9.5e-4 |
| Readout threshold | 0.27 | 0.17 | 0.75 |
| Gain strength | 0.20 | 0.057 | 0.11 |

**Figure S4:**
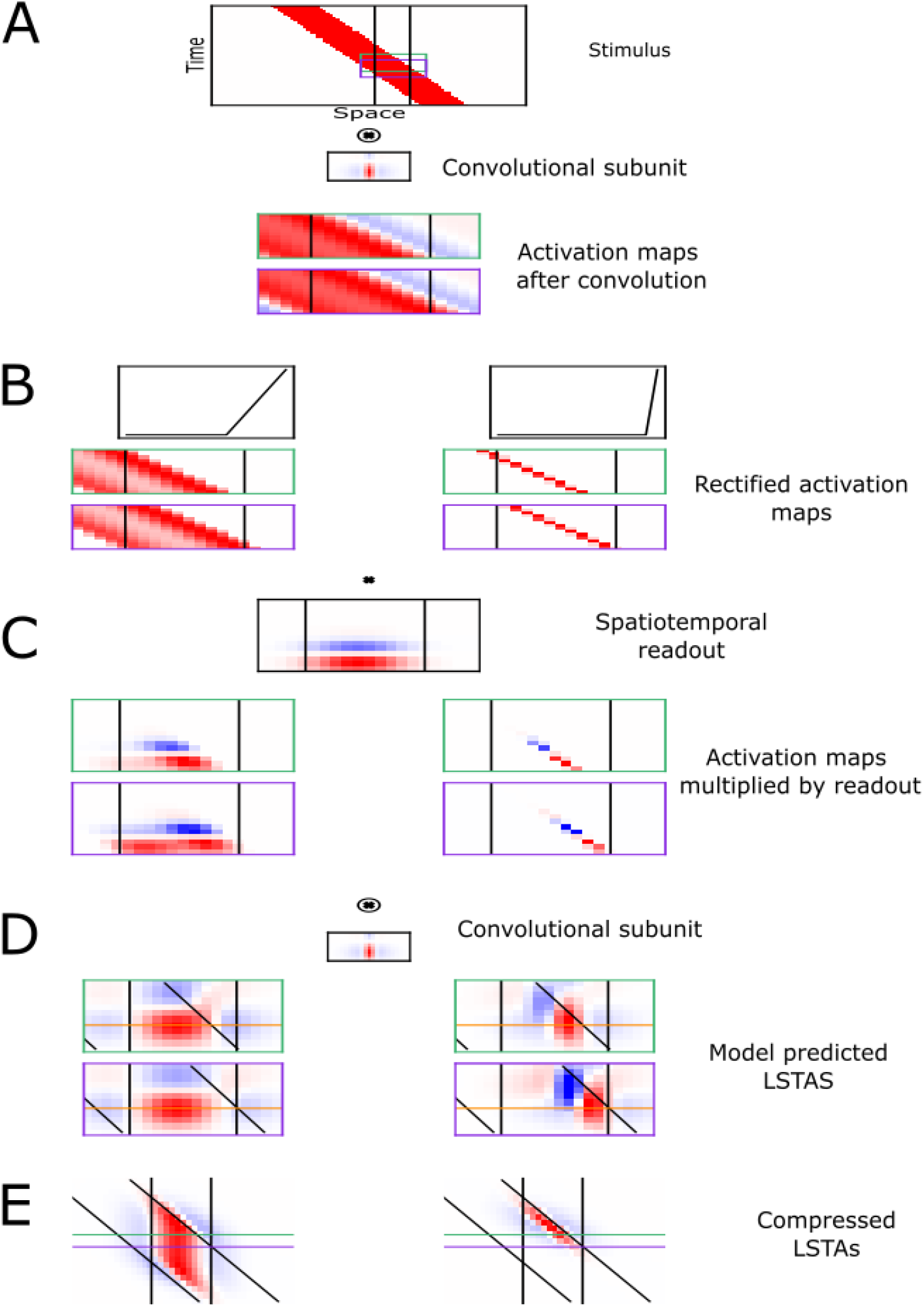
A simplified subunit model demonstrates how increasing the rectification threshold shifts model LSTAs from static to edge-tracking. To directly test the effect of subunit rectification threshold on LSTA behavior, we constructed a simplified version of our model. The stimulus was processed by a single ON subunit (identical to that used in the full model), and the spatiotemporal readout filter was derived by denoising the modeled cell’s classical STA. The readout nonlinearity was removed (threshold set to zero), and no gain control was included. The subunit rectification threshold was the only free parameter. **A**. Top: bar stimulus in space-time coordinates. The two colored boxes indicate successive time windows at which LSTAs are computed. Vertical bars mark the spatial boundaries of the cell’s classical RF. Middle: the ON convolutional subunit. Bottom: The two activation maps obtained by convolving the stimulus within each time window with the ON bipolar subunit. **B**. Subunit activation maps after applying low (left) and high (right) rectification thresholds. The nonlinear rectification functions applied to the subunit outputs are shown above each condition. At low threshold, subunit responses are spatially broad; at high threshold, activity becomes localized to the bar’s leading edge. **C**. Top: Spatiotemporal readout for an example cell. Bottom: element-wise product of the rectified activation maps with the readout filter. To obtain the model’s predicted firing rate, this map is summed over space and time. To compute the model-predicted LSTA, this map is instead deconvolved with the subunit filter (model gradient). **D**. Top: the subunit is shown again to indicate its use in the deconvolution step. Bottom: model-predicted LSTAs for the two time windows under low (left) and high (right) threshold conditions. **E**. Compressed LSTA representations (as in main figures), illustrating the transition from static selectivity (left) to edge-tracking (right) as threshold increases. Vertical black lines denote the classical RF, diagonal lines denote bar edges, and horizontal colored lines indicate 1D LSTAs slices taken at the example time windows.

**Video V1:**
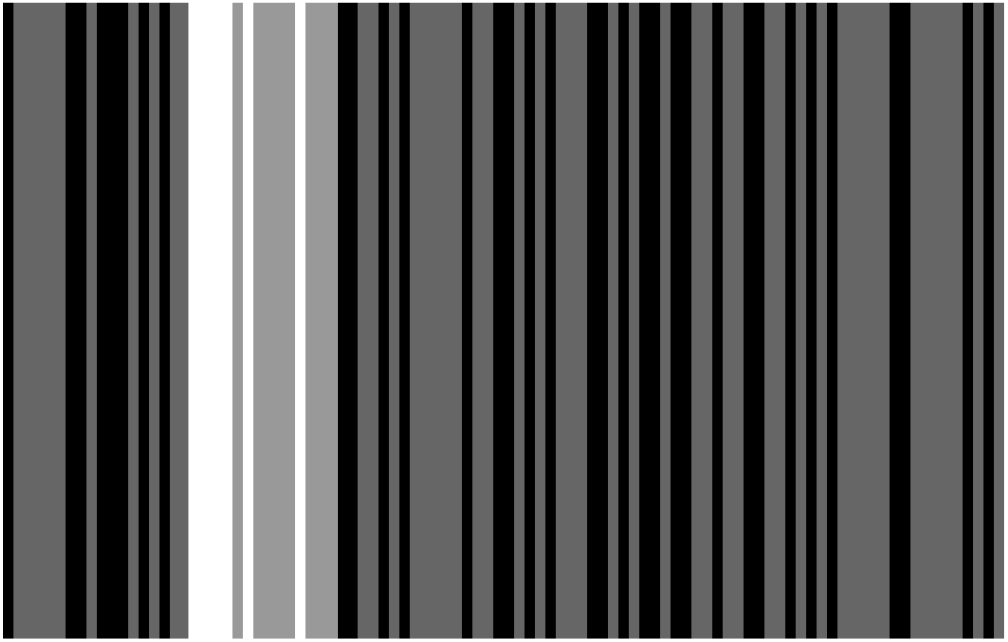
Example of a perturbed bar sweep played at half speed for ease of visual inspection. The full recording consisted of 2050 of such sweeps, each time superimposed with a different random perturbation. See Methods for stimulus details.

**Video V2:**
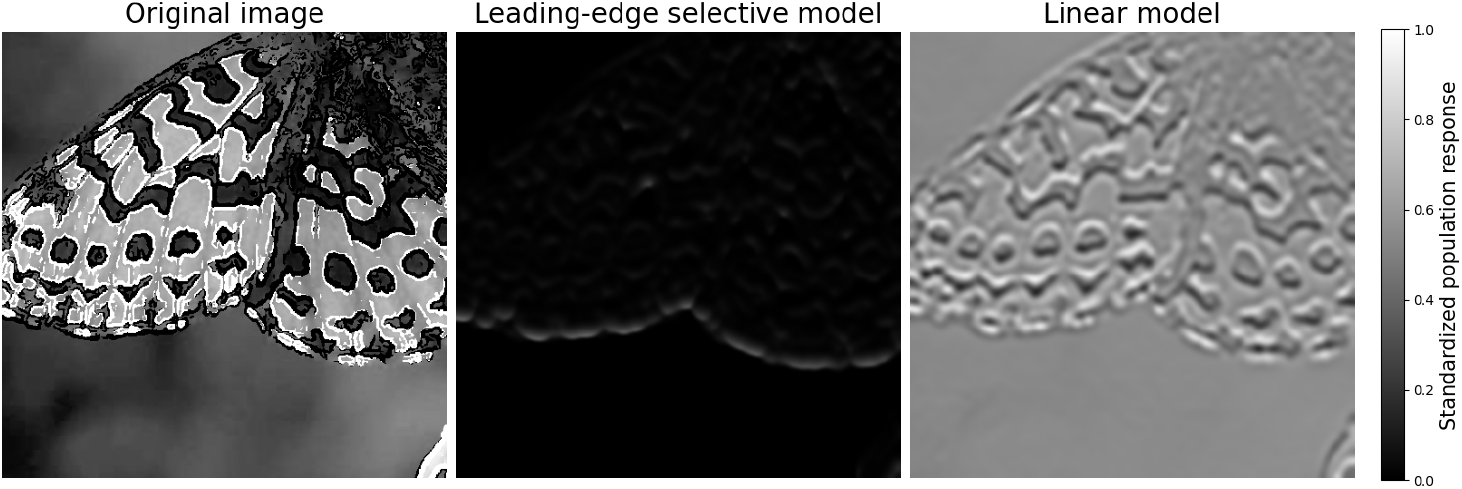
Dynamic extraction of moving textured edges in a naturalistic stimulus. The video shows a downward-drifting butterfly stimulus (left), alongside the responses of two computational models. The center panel displays the output of the leading-edge selective model, in which the stimulus is first convolved with spatiotemporal difference-of-Gaussians subunits, rectified, convolved with a larger Gaussian filter to simulate a population of ganglion cells, and modulated by divisive gain control. The right panel shows the output of a corresponding linear model that omits rectification and gain control while preserving the filtering stages. Model responses are standardized to the range [0,1] for visualization. All the parameters of the models were chosen by hand in arbitrary units. The purpose of these models is purely illustrative.

## Notes

### Competing Interest Statement

The authors have declared no competing interest.

### Summary of Updates

One illustrative figure was added in the discussion.

## References

1. Pasupathy, A. THE NEURAL BASIS OF IMAGE SEGMENTATION IN THE PRIMATE BRAIN. Neuroscience 296, 101–109 (2015).

2. Kuffler, S. W. Discharge patterns and functional organization of mammalian retina. J. Neurophysiol. 16, 37–68 (1953).

3. Wienbar, S. & Schwartz, G. W. The dynamic receptive fields of retinal ganglion cells. Prog. Retin. Eye Res. 67, 102–117 (2018).

4. Marr, D. & Hildreth, E. Theory of edge detection. Proc. R. Soc. Lond. B Biol. Sci. 207, 187–217 (1980).

5. Turner, M. H. & Rieke, F. Synaptic rectification controls nonlinear spatial integration of natural visual inputs. Neuron 90, 1257–1271 (2016).

6. Barlow, H. B. & Hill, R. M. Selective sensitivity to direction of movement in ganglion cells of the rabbit retina. Science 139, 412–414 (1963).

7. Wei, W. Neural Mechanisms of Motion Processing in the Mammalian Retina. Annu. Rev. Vis. Sci. 4, 165–192 (2018).

8. Berry, M. J., Brivanlou, I. H., Jordan, T. A. & Meister, M. Anticipation of moving stimuli by the retina. Nature 398, 334–338 (1999).

9. Johnston, J. & Lagnado, L. General features of the retinal connectome determine the computation of motion anticipation. eLife 4, e06250 (2015).

10. DePiero, V. J. & Borghuis, B. G. Phase Advancing Is a Common Property of Multiple Neuron Classes in the Mouse Retina. eNeuro 9, (2022).

11. Manookin, M. B., Patterson, S. S. & Linehan, C. M. Neural Mechanisms Mediating Motion Sensitivity in Parasol Ganglion Cells of the Primate Retina. Neuron 97, 1327–1340.e4 (2018).

12. Gaynes, J. A., Budoff, S. A., Grybko, M. J., Hunt, J. B. & Poleg-Polsky, A. Classical center-surround receptive fields facilitate novel object detection in retinal bipolar cells. Nat. Commun. 13, 5575 (2022).

13. Baden, T. et al. The functional diversity of retinal ganglion cells in the mouse. Nature 529, 345–350 (2016).

14. Goldin, M. A. et al. Context-dependent selectivity to natural images in the retina. Nat. Commun. 13, 5556 (2022).

15. Chichilnisky, E. J. A simple white noise analysis of neuronal light responses. Netw. Bristol Engl. 12, 199–213 (2001).

16. Enroth-Cugell, C. & Robson, J. G. The contrast sensitivity of retinal ganglion cells of the cat. J. Physiol. 187, 517–552 (1966).

17. Gollisch, T. & Meister, M. Rapid neural coding in the retina with relative spike latencies. Science 319, 1108–1111 (2008).

18. Deny, S. et al. Multiplexed computations in retinal ganglion cells of a single type. Nat. Commun. 8, 1964 (2017).

19. Chen, E. Y. et al. Alert Response to Motion Onset in the Retina. J. Neurosci. 33, 120–132 (2013).

20. Chen, E. Y., Chou, J., Park, J., Schwartz, G. & Berry, M. J. The Neural Circuit Mechanisms Underlying the Retinal Response to Motion Reversal. J. Neurosci. 34, 15557–15575 (2014).

21. Schwartz, G. W. et al. The spatial structure of a nonlinear receptive field. Nat. Neurosci. 15, 1572–1580 (2012).

22. Strauss, S. et al. Center-surround interactions underlie bipolar cell motion sensitivity in the mouse retina. Nat. Commun. 13, 5574 (2022).

23. Gollisch, T. & Meister, M. Eye Smarter than Scientists Believed: Neural Computations in Circuits of the Retina. Neuron 65, 150–164 (2010).

24. Leonardo, A. & Meister, M. Nonlinear Dynamics Support a Linear Population Code in a Retinal Target-Tracking Circuit. J. Neurosci. 33, 16971–16982 (2013).

25. Schwartz, G., Taylor, S., Fisher, C., Harris, R. & Berry, M. J. Synchronized Firing among Retinal Ganglion Cells Signals Motion Reversal. Neuron 55, 958–969 (2007).

26. Marre, O. et al. High Accuracy Decoding of Dynamical Motion from a Large Retinal Population. PLOS Comput. Biol. 11, e1004304 (2015).

27. Karamanlis, D. & Gollisch, T. Nonlinear Spatial Integration Underlies the Diversity of Retinal Ganglion Cell Responses to Natural Images. J. Neurosci. 41, 3479–3498 (2021).

28. Yu, Y., Stirman, J. N., Dorsett, C. R. & Smith, S. L. Selective representations of texture and motion in mouse higher visual areas. Curr. Biol. 32, 2810–2820.e5 (2022).

29. Angueyra, J. M., Baudin, J., Schwartz, G. W. & Rieke, F. Predicting and Manipulating Cone Responses to Naturalistic Inputs. J. Neurosci. 42, 1254–1274 (2022).

30. Demb, J. B., Zaghloul, K. & Sterling, P. Cellular Basis for the Response to Second-Order Motion Cues in Y Retinal Ganglion Cells. Neuron 32, 711–721 (2001).

31. Borghuis, B. G., Marvin, J. S., Looger, L. L. & Demb, J. B. Two-Photon Imaging of Nonlinear Glutamate Release Dynamics at Bipolar Cell Synapses in the Mouse Retina. J. Neurosci. 33, 10972–10985 (2013).

32. Trenholm, S. et al. Nonlinear dendritic integration of electrical and chemical synaptic inputs drives fine-scale correlations. Nat. Neurosci. 17, 1759–1766 (2014).

33. Drinnenberg, A. et al. How Diverse Retinal Functions Arise from Feedback at the First Visual Synapse. Neuron 99, 117–134.e11 (2018).

34. Zoccolan, D., Oertelt, N., DiCarlo, J. J. & Cox, D. D. A rodent model for the study of invariant visual object recognition. Proc. Natl. Acad. Sci. U. S. A. 106, 8748–8753 (2009).

35. Froudarakis, E. et al. Object manifold geometry across the mouse cortical visual hierarchy. 2020.08.20.258798 Preprint at 10.1101/2020.08.20.258798 (2021).

36. Siegenthaler, D. et al. Visual objects refine head direction coding. Science 389, eadu9828 (2025).

37. Hoy, J. L., Yavorska, I., Wehr, M. & Niell, C. M. Vision Drives Accurate Approach Behavior during Prey Capture in Laboratory Mice. Curr. Biol. 26, 3046–3052 (2016).

38. Luongo, F. J. et al. Mice and primates use distinct strategies for visual segmentation. eLife 12, e74394 (2023).

39. Feldman, J. What is a visual object? Trends Cogn. Sci. 7, 252–256 (2003).

40. Mishkin, M., Ungerleider, L. G. & Macko, K. A. Object vision and spatial vision: two cortical pathways. Trends Neurosci. 6, 414–417 (1983).

41. DiCarlo, J. J., Zoccolan, D. & Rust, N. C. How Does the Brain Solve Visual Object Recognition? Neuron 73, 415–434 (2012).

42. Marre, O. et al. Mapping a Complete Neural Population in the Retina. J. Neurosci. 32, 14859–14873 (2012).

43. Yger, P. et al. A spike sorting toolbox for up to thousands of electrodes validated with ground truth recordings in vitro and in vivo. eLife 7, e34518 (2018).

44. Park, M., Horwitz, G. & Pillow, J. Active learning of neural response functions with Gaussian processes. in Advances in Neural Information Processing Systems vol. 24 (Curran Associates, Inc., 2011).

45. Goldin, M. A., Virgili, S. & Chalk, M. Scalable Gaussian process inference of neural responses to natural images. Proc. Natl. Acad. Sci. 120, e2301150120 (2023).

46. Duncker, L., Ruda, K. M., Field, G. D. & Pillow, J. W. Scalable Variational Inference for Low-Rank Spatiotemporal Receptive Fields. Neural Comput. 35, 995–1027 (2023).

